# GalaxyMemMD: an end-to-end Galaxy toolset for reproducible membrane embedding, molecular dynamics simulation, and analysis of membrane proteins

**DOI:** 10.64898/2026.09.28.755125

**Authors:** Jayadev Joshi, Bryan Raubenolt, Natalie Katzenmeyer, Fabio Cumbo, Daniel Blankenberg

## Abstract

Membrane proteins play crucial roles in cellular signaling, transport, and pharmacological regulation, yet their structural and functional characterization remains challenging due to experimental limitations. Molecular dynamics (MD) simulations have become indispensable for probing membrane protein behavior at atomic resolution; however, executing these simulations at scale remains a significant bottleneck. Existing tools excel at system preparation for individual cases, but none offer a complete, high-throughput pipeline spanning embedding, simulation, and analysis across multiple membrane compositions, and rarely integrate modern structure prediction and molecular docking frameworks.

Here, we present GalaxyMemMD, a fully automated, open-source Galaxy framework that provides a direct, scalable alternative to existing preparation-only tools. GalaxyMemMD integrates every stage membrane embedding, system preparation, MD execution, and downstream analysis into a single reproducible pipeline. Leveraging Galaxy’s HPC-ready job scheduling, GalaxyMemMD natively supports high-throughput parallel simulations across multiple membrane compositions and protein variants, including AlphaFold-derived structural inputs and downstream molecular docking workflows, without repeated manual setup.

We validate GalaxyMemMD across six structurally diverse membrane proteins four GPCRs, one ion channel, and one beta-barrel demonstrating consistent system preparation, while the benchmark analysis shows stable embeddings, and reproducible trajectory-level production workflow. Across these benchmarks, GalaxyMemMD matches the setup quality of established tools, therefore serves as both a practical alternative and a capability extension for the membrane protein simulation community.

## 1. Introduction

Membrane proteins constitute nearly 30% of the human proteome and are targeted by more than 50% of approved therapeutic drugs, highlighting their central biomedical importance ^11,2^ Among the best-characterized membrane proteins are G protein–coupled receptors (GPCRs), the largest membrane protein family in the human genome and master regulators of nearly every major physiological process, including neurotransmission, hormone signaling, and sensory perception. The biomedical significance of GPCRs is unparalleled: approximately one-third of all FDA-approved drugs exert their effects through GPCR targets, underscoring their central role in disease and therapeutic intervention ^3^. Despite their significance, membrane proteins remain exceptionally challenging to study experimentally ^4^. Their hydrophobic transmembrane regions, dependence on native lipid environments for structural integrity, and intrinsic conformational flexibility make purification, crystallization, and structural characterization difficult ^5,6^. Even with advances in cryo-electron microscopy (cryo-EM), NMR spectroscopy, and lipid-bilayer mimetics such as nanodiscs, obtaining high-resolution GPCR structures that preserve native-like lipid–protein interactions remains technically challenging and often requires system-specific optimization of receptor stabilization, membrane reconstitution, and experimental conditions ^6,7^. Yet even when high-resolution structures are obtained, they provide only a static snapshot: crystal packing artifacts, cryo-cooling, and detergent solubilization can distort native conformations, and the inherently dynamic nature of membrane protein function conformational switching, lipid-coupled motions, and ligand-induced allosteric rearrangements remains invisible to static structural methods. These persistent experimental limitations have made computational modeling and MD simulations indispensable, as they enable atomic-level exploration of membrane protein dynamics, conformational transitions, and lipid interactions that are difficult or impossible to capture experimentally ^8,9^.

Several computational platforms such as GROMACS ^4,10^, AMBER ^11^, CHARMM ^12,13^ and NAMD ^14^ have been developed to facilitate MD simulation of proteins. However, simulating membrane proteins still remains challenging due to its complex system preparation, size, and component heterogeneity. Tools such as CHARMM-GUI ^15–17^, GROMACS web servers, and MDWeb provide user-friendly interfaces for system preparation, solvation, ion addition, and initial simulation setup ^18^. However, these tools focus primarily on system preparation and do not offer an integrated, end-to-end environment that links setup, production simulation, and trajectory analysis in a single reproducible workflow.

CHARMM-GUI, the most widely used preparation tool, stops at system setup and provides production ready system topologies to extend the MD simulation up to the desired time period. However, it requires manual simulation management in a separate environment beyond the production ready topologies. MDWeb and VMD-based approaches similarly lack automated MD execution, multi-membrane batch workflows, and native integration with structure prediction tools such as AlphaFold or downstream docking protocols ^19,20^. Researchers who need to systematically compare membrane compositions, screen protein variants, or run large-scale simulation campaigns must therefore invest substantial effort entirely outside these tools. There is a clear and unmet need for a reproducible, scalable, and fully automated framework that serves as a high-throughput alternative one that spans structure input through post-MD analysis and leverages HPC infrastructure for parallel execution across membrane environments.

To address these unmet needs, we developed GalaxyMemMD, an open-source, reproducible, and fully automated Galaxy-based alternative to existing tools, offering a complete, high-throughput-capable workflow-based framework for protein-membrane simulations, built on the Galaxy platform ^21^. Galaxy’s ecosystem inherently supports transparency, version control, and FAIR computational practices, enabling users to share complete, executable workflows and histories with full provenance information. Leveraging these strengths, GalaxyMemMD integrates each stage of membrane protein modeling from membrane embedding to system preparation, MD simulation, and downstream structural analysis into a cohesive, modular pipeline that minimizes manual intervention and ensures methodological consistency across studies. A key objective of GalaxyMemMD is to make advanced membrane simulations accessible to researchers with diverse computational backgrounds, while still providing the scalability required for expert-level analyses. This uniquely supports comparative studies of lipid environments, an analysis that is technically demanding with existing tools and often requires extensive scripting or manual system setup. Importantly, GalaxyMemMD is designed to be generalizable across protein classes, including GPCRs, multi-pass transporters, ion channels, enzymes, and other integral membrane proteins protein families that are increasingly being studied using MD owing to advances in structural prediction tools such as AlphaFold ^19^ and RoseTTAFold ^22^. By natively integrating these prediction frameworks within the workflow, GalaxyMemMD simplifies the overall MD analysis of these complex proteins. We further validate our method by applying it to 6 diverse membrane proteins: 4 G-protein coupled receptors (GPCRs), 1 beta-barrel, and 1 ion channel (see Figure 1 below). Together, these capabilities position GalaxyMemMD as a robust, high-throughput-ready alternative to the existing tools, and lowers the barrier to large-scale protein-membrane simulations while enhancing reproducibility, methodological transparency, and interoperability across the broader computational biology community.

**Figure 1.**
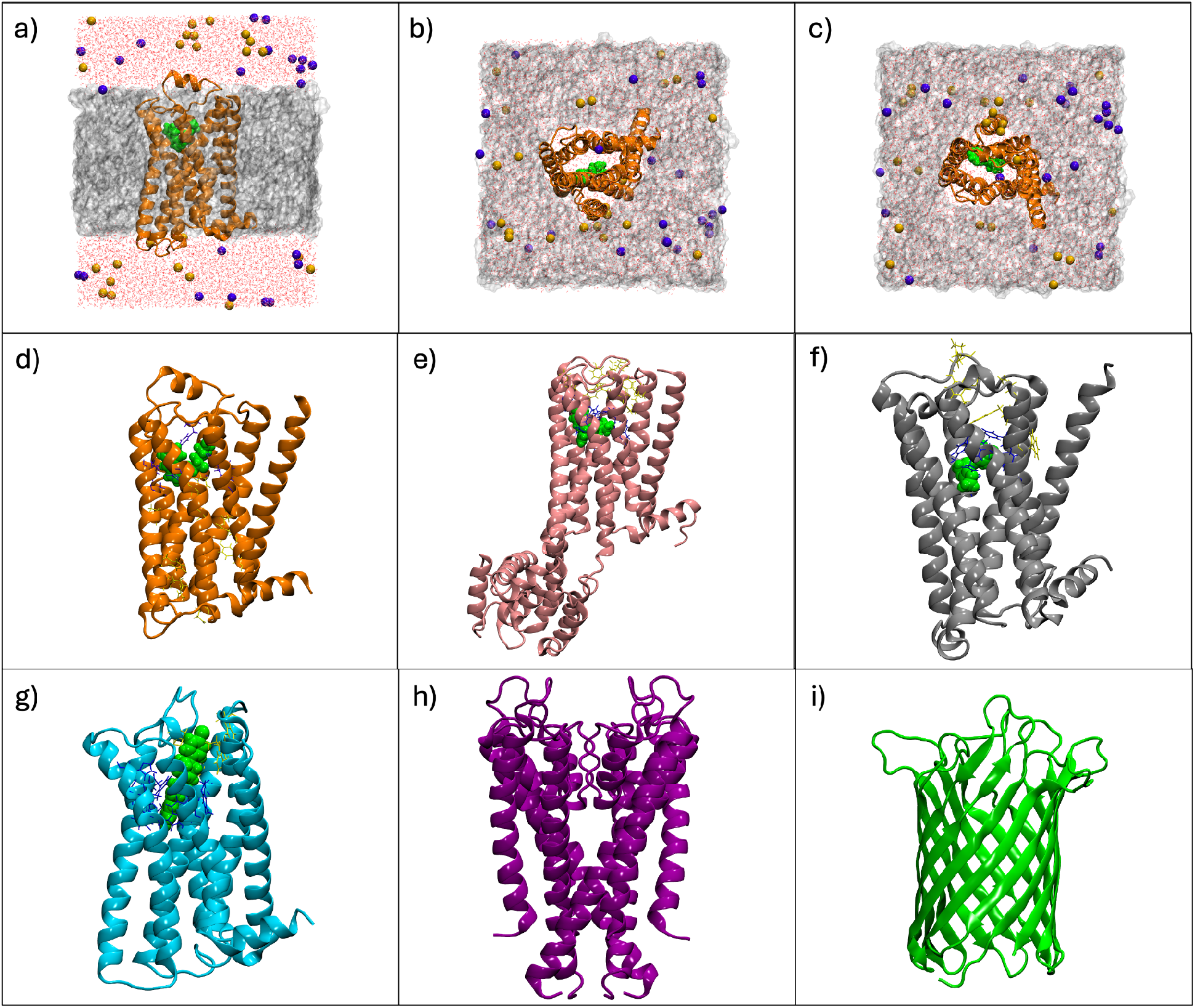
An example of a final built system seen from a) a lateral view (membrane slice), b) outward from the cytosol, c) inward from the extracellular media. In all cases, the water molecules are red lines, the K+Cl-ions are purple and yellow, the drug/ligand in green, and the lipid bilayer in grey. The 6 modeled protein-membrane systems include the d) β2-adrenergic receptor (PDB: 2RH1) ^23^, e) M1 muscarinic receptor (PDB: 5CXV) ^24^, f) M2 muscarinic receptor (PDB: 4MQS) ^25^, g) D2 dopamine receptor (PDB: 6LUQ) ^26^, h) potassium ion channel (PDB: 1R3J), and i) the beta barrel Ompg (PDB: 2IWW) ^27^. For the complexes in d-g, the orthosteric residues are colored in blue, while the allosteric residues are in yellow.

## 2. Materials and Methods

GalaxyMemMD was built upon the open-source, web-based Galaxy platform that enables accessible and fully reproducible biomedical data analysis through a unified graphical environment^21^. It provides versioned and dependency-isolated tool execution, automated provenance tracking of all parameters and datasets, and scalable workflow execution across local, HPC, and cloud infrastructures. Tools and workflows are curated through the Galaxy ToolShed ^28^, supporting community-driven method sharing and adherence to FAIR principles. Together, these features make Galaxy a robust and extensible framework for transparent, standardized, and reproducible computational research. A detailed architectural overview is as follows.

### 2.1 Comparison With Existing Tools

Several established platforms support protein-membrane system preparation and molecular simulation setup, including CHARMM-GUI ^15,16^, MDWeb ^18^, VMD membrane-building ^29^ plugins, and stand-alone PACKMOL/MemGen or AmberTools scripts. Each of these resources has made important contributions to advancing accessibility in computational structural biology. However, they typically operate as single-purpose preparation interfaces or require manual scripting, whereas the Galaxy-based framework presented here offers a unified, automated, and reproducible end-to-end system that integrates structure acquisition, membrane construction, solvation and ionization, topology generation, simulation execution, solvent-structure modeling, water placement, analysis, and downstream docking integration. Below, we summarize conceptual and functional distinctions between our workflow and current alternatives.

The Galaxy-based workflow provides a modular HPC-ready system capable of executing multiple membrane compositions, simulations, and analyses in parallel. Crucially, Galaxy’s FAIR-compliant reproducibility engine records all parameters, tool versions, and dataset histories, enabling transparent and shareable workflows capabilities largely absent from other web-based or script-driven alternatives.

Table 1 shows a comprehensive feature comparison from the existing tools. CHARMM-GUI generates inputs for multiple MD engines, but does not manage the data and end-to-end execution. MDWeb/MDMoby is for composable workflows and analysis, but does not do production execution because only test simulations run server-side. VMD enables local deployment, interoperability, and analysis, but does not offer a complete workflow engine or a full-platform open-source license because the VMD core and plugin licenses differ. The composite comparison in Table 1 illustrates the feature presence of the selected platforms rather than ranking platform quality or performance. The platforms differ in scope and deployment model: some primarily generate systems or scripts, some provide modular services or local plugins, and GalaxyMemMD emphasizes workflow-managed execution, provenance, and analysis.

**Table 1.** Feature-presence comparison of selected protein–membrane simulation platforms. Statuses follow the operational definitions and dated source review provided below.

| Criterion | CHARMM-GUI<sup>15</sup> | MDWeb/MD Moby<sup>18</sup> | VMD membrane tools<sup>29</sup> | GalaxyMemMD |
| --- | --- | --- | --- | --- |
| **Complete end-to-end workflow** | No | No | No | Yes |
| **Input and Output data management** | No | No | No | Yes |
| **Local/self-hosted deployment** | No | No | Yes | Yes |
| **Composable workflow engine** | No | Yes | No | Yes |
| **Versioned provenance and workflow export** | No | No | No | Yes |
| **Fully open-source platform** | No | No | No | Yes |
| **Interoperability with other tools/engines** | Yes | Yes | Yes | Yes |
| **Post-MD trajectory analysis** | No | Yes | Yes | Yes |
| **Status definitions. Yes:** the feature is provided within the evaluated platform scope according to the cited source and limited testing. **No:** the feature is not provided within that scope. |  |  |  |  |

#### Scope and sources

This is a documentation and published research article based feature comparison, we do not perform a full scale benchmark against all the listed tools except CHAMM-GUI. Documentation was reviewed on 26 September 2026. Sources: CHARMM-GUI Membrane Builder documentation and [FAQ](https://www.charmm-gui.org/?doc=faq); [MDWeb/MDMoby publication](https://academic.oup.com/bioinformatics/article/28/9/1278/313437); [VMD plugin documentation](https://www.ks.uiuc.edu/Research/vmd/plugins/) and [license](https://www.ks.uiuc.edu/Research/cloud/LICENSE.html); [Galaxy reproducibility documentation](https://training.galaxyproject.org/training-material/topics/introduction/tutorials/galaxy-reproduce/tutorial.html) and [job-configuration documentation](https://docs.galaxyproject.org/en/release_18.05/admin/jobs.html). GalaxyMemMD entries are based on this manuscript and should be linked to exact ToolShed/workflow versions in Data Availability.

### 2.2 Architectural Overview of the GalaxyMemMD

Tool integration in Galaxy is facilitated through XML-based tool wrappers, which define command execution logic, parameter handling, data types, and post-processing steps. GalaxyMemMD leverages this framework to encapsulate each of its 11 modular tools (Table 2), ensuring consistent behavior, standardized I/O handling, and compatibility with Galaxy’s workflow editor. These wrappers allow researchers to customize individual steps, for example, changing force-field parameters, simulation lengths, or membrane compositions while maintaining an end-to-end automated pipeline structure. These newly developed tools, as well as the legacy computational chemistry tools used in the full workflows are available in the Galaxy ToolShed and the repositories for workflows can be found in the data availability section.

**Table 2.** GalaxyMemMD toolset. This toolset comprises 11 tools spanning system preparation, simulation setup and conversion, and water modeling.

| **Tool name** | **Functionality** | **Major Libraries and tools uses** |
| --- | --- | --- |
| *AMBER compatible PDB converter* | Convert PDB files to AMBER-compatible format | Python, biobb_amber, XML tool wrapper |
| *AMBER to GROMACS* | Converts AMBER topology and coordinate files to GROMACS format | Python, ParmEd, XML tool wrapper |
| *Generate GROMACS MDP File* | Generate a GROMACS .mdp file with presets and advanced controls | Python, GROMACS, XML tool wrapper |
| *Generate custom GROMACS index* | Generate GROMACS index files with custom groups for analysis and simulation | Python, GROMACS, XML tool wrapper |
| *Membrane Builder* | Embed proteins into lipid bilayer membranes for MD simulations | AmberTools, Python, XML tool wrapper |
| *Prepare AMBER System for solvation* | Build AMBER parameter files and define a box to predict the solvent around protein | Python, AmberTools, numpy, XML tool wrapper |
| *Shift Membrane* | Fix membrane position in respect to the protein | Python, XML tool wrapper |
| *Solvent Laplacian Calculator* | Calculate laplacian for solvent around protein | Python, AmberTools, XML tool wrapper |
| *Solvent Density Calculator* | Calculate solvent density around protein | Python, AmberTools, XML tool wrapper |
| *Place Solvent Molecules* | Place solvent around protein | Python, AmberTools, XML tool wrapper |
| *Merge Water Residues* | Extract selected water residues and merge into a system PDB file | Python, XML tool wrapper |

Together, these architectural features make Galaxy an ideal backbone for GalaxyMemMD, enabling scalable, reproducible, and user-friendly deployment of sophisticated protein-membrane simulation workflows.

#### 2.2.1 Membrane-embedding and Simulation Tools

a. **AMBER compatible PDB converter tool** prepares PDB files for MD simulation using PDB4AMBER which converts Protein Data Bank (PDB) files into an AMBER-compatible format suitable for simulation with the AMBER force fields. During this step, atom-naming conventions were standardized and common PDB formatting inconsistencies were resolved. Hydrogen-bond networks were optimized with Reduce, and alternate conformations were resolved to single occupancies. Non-standard residues were retained or removed as appropriate for the system, and water molecules and ions were managed to yield a clean, force-field-consistent starting structure.
b. **AMBER to GROMACS tool** *converts* parameterized AMBER topologies to their GROMACS equivalents using an AMBER-to-GROMACS conversion step. The AMBER topology (prmtop/parm7) and coordinate (inpcrd/rst7) files were read and converted into a GROMACS topology (.top) and structure (.gro) file, preserving the force-field parameters, atom types, bonded and non-bonded terms, and charges defined under AMBER. This allowed the AMBER-parameterized system, including the explicitly placed internal water molecules to be propagated in GROMACS while retaining the original force-field description.
c. **Generate GROMACS MDP File tool** generates simulation control parameters. Since the .mdp file governs every aspect of a GROMACS run its correct and internally consistent specification is essential yet error-prone when edited by hand, and is a common source of irreproducibility across studies. This tool addresses this by generating validated parameter files in two modes. In workflow mode, it produces a complete, staged equilibration-to-production protocol for membrane-embedded systems two NVT stages for thermal stabilization at constant volume, followed by four successive NPT stages with progressively released position restraints, and an unrestrained production run together with the corresponding GROMACS index file (.ndx) and a human-readable list of atom groups generated directly from the system structure. Group definitions are drawn from the membrane-builder configuration (lipids, ligand, ions), ensuring that the temperature-coupling and centre-of-mass groups referenced in the .mdp files are consistent with the index file and thereby eliminating a frequent cause of run failure. In custom mode, the tool exposes the full parameter set so that advanced protocols remain fully configurable. By encapsulating a standardized, membrane-aware equilibration protocol behind a graphical interface and embedding it in a reproducible Galaxy workflow, this tool lowers the technical barrier to correctly configured simulations, reduces manual-editing errors, and makes the exact simulation parameters transparent and shareable an important consideration for the reproducibility of the overall pipeline.
d. **Generate custom GROMACS index tool** generates atom-group definitions for simulation control and analysis. The index file (.ndx) specifies the atom selections used for temperature and pressure coupling, position restraints, output control, and analysis. From the system structure (.gro) and a configuration file enumerating the molecular components (protein, individual lipid types, ligand, and ions), the tool builds the corresponding index groups automatically, including standard, per-component, and user-defined combined selections. Because these definitions derive from the same component configuration used elsewhere in the pipeline, the index file stays consistent with the groups referenced in the simulation parameter files, removing a common source of run failure and supporting a reproducible setup.
e. **Membrane Builder tool** provides an automated and customizable framework for constructing protein-membrane systems suitable for molecular dynamics simulations. This tool is implemented using the python-based Packmol-Memgen package ^30^ and AmberTools (v24.8) ^31^, which serve as its primary dependencies. Given a user-supplied protein structure (PDB), lipid composition and mixture ratios, the tool embeds the protein into one or more lipid bilayers, supporting asymmetric leaflets, and can generate a solvated system with user-defined water-layer thickness, and box dimensions. While default settings enable rapid, standardized model construction, advanced options offer fine-grained control during the membrane embedding step. The tool’s inputs therefore range from minimal (protein, lipids, ratios, geometric distances) to fully configured multi-parameter builds, while its outputs consist of a complete, simulation-ready embedded protein-membrane system (PDB) and an accompanying log file (TXT) capturing all executed steps for reproducibility. Together, these capabilities allow the *Membrane Builder tool* to generate complex, biologically realistic membrane environments while maintaining transparency, flexibility, and workflow-level reproducibility within the Galaxy framework. This tool produces a system PDB file and a configuration file, which help in generating the MDP files automatically in the next step. In addition, it can optionally generate the ligand file separately.
f. **Shift Membrane tool** provides an automated method to realign protein-membrane systems by translating the membrane-embedded model so that the protein occupies the correct spatial position relative to its original, unmodified structure. The tool calculates the center of mass (COM) of both the original protein and the protein-membrane complex excluding lipid, solvent, and ion atoms and computes the required translation vector to reposition the membrane so that the embedded protein matches the orientation of the reference protein. This correction step is often required after membrane building procedures in order to restore consistent coordinate frames before downstream simulation or analysis. Users provide two PDB files: (i) the original protein structure and (ii) the membrane-embedded protein generated by Packmol-Memgen or related builders. The tool outputs a shifted protein-membrane system (PDB) with corrected coordinates, preserving all membrane, solvent, and ion entries while applying a uniform translation based on COM alignment. Additionally, it reports the system’s approximate box dimensions computed from the solvent distribution, allowing users to verify spatial consistency. This tool ensures that protein-membrane systems maintain appropriate alignment and coordinate integrity as part of a reproducible Galaxy-based MD workflow.

#### 2.2.2 Tools for 3D-RISM- and Laplacian-Guided Placement of Water Molecules

Accurate placement of internal and interfacial water molecules is important across membrane proteins because buried waters can stabilize structure, bridge polar or charged residues, mediate ligand or cofactor binding, participate in proton or ion transport, and influence conformational transitions. In membrane proteins, functionally important waters may occupy deeply buried, occluded cavities whose exchange with bulk solvent is far slower than accessible equilibration MD-radiolytic-labeling experiments detect essentially no exchange of the core structural waters with bulk solvent across functional states ^32^, so naïve box solvation followed by standard equilibration frequently leaves these cavities empty or mis-hydrated, biasing the membrane protein toward artifactual conformations and distorting the internal hydrogen-bond network ^33^. GPCRs provide one well-studied example: accurate hydration of the transmembrane core is a prerequisite for meaningful GPCR simulations, because water in these receptors is not a passive solvent but an integral structural and functional element ^34,35^. Similar considerations apply to other membrane-bound receptors, channels, transporters, enzymes, and complexes whenever buried hydration contributes to stability or mechanism. If analysis is sensitive to internal hydration, our implemented workflow provides an option for users to explicitly reconstruct those missing water molecules before full system simulation ^36,37^.

a. **Prepare AMBER System for solvation tool** prepares protein or protein–ligand structures for subsequent solvent prediction. The tool accepts a protein structure in PDB format and optionally ligand coordinates and parameters in MOL2 and FRCMOD formats. The simulation bounding box can either be calculated automatically from the van der Waals extent of the cleaned protein with a user-defined buffer (10 Å by default) or specified manually using X, Y, and Z dimensions and unit-cell angles. The tool outputs an AMBER topology (PARM7), a boxed coordinate/restart file (RST7), the tleap-generated system PDB, and the cleaned protein-only PDB, providing the required system files for downstream solvent prediction.
b. **Solvent Density Calculator tool** is implemented based on the AmberTools, estimates solvent distribution around biomolecules by solving integral equations that describe solute–solvent interactions, generating volumetric maps that characterize hydration structure, ion localization, and solvent thermodynamics. Users supply the required AMBER input files, namely the RISM control file (mdin), parameter/topology file (parm7), coordinate/restart file (rst7), and optionally an external solvent susceptibility function (.xvv) and the tool executes Sander in 3D-RISM mode is used. The tool produces several outputs: the primary 3D-RISM results file (out.kh.r3d), DX-format volumetric maps for hydrogen and oxygen interaction sites (out.kh.H1.0.dx and out.kh.O.0.dx), and an mdinfo log summarizing run parameters and calculation progress calculated based on cSPCE_kh.xvv water model. This module enables reproducible solvent-structure analysis and facilitates downstream applications such as implicit hydration modeling, pocket detection, and membrane system refinement.
c. **Solvent Laplacian Calculator tool** performs Laplacian-based analysis of volumetric density maps generated from 3D-RISM calculations, enabling the identification of curvature features and localized density variations within solvent distributions. Using AmberTools and NumPy (v1.23.3) as dependencies, METATWIST processes an input DX file, typically the oxygen-site density map (kh.O.0.dx) to compute its Laplacian and optionally apply Gaussian convolution controlled by user-defined parameters. The tool accepts the density map along with the species identifier, convolution mode, and sigma value, and produces two key outputs: (i) a Laplacian profile text file summarizing density curvature information and (ii) an updated DX file (out.kh.O.dx) containing the processed density field. These outputs are essential for downstream analysis such as solvent-feature detection, hydration-site characterization, and preparing enhanced density maps used by water-placement or cavity-detection tools.
d. **Placement of water molecules tool** automates the identification and placement of structurally relevant water molecules within a protein environment by analyzing volumetric density maps generated 3D-RISM calculation using AmberTools and NumPy as dependencies. The tool processes two electrostatic potential–derived DX files typically kh.O.0.dx and kh.O.dx generated by Sander to detect high-probability hydration sites. Users provide the two DX files along with parameters controlling the density map interpretation (species identifier, bulk density, and detection threshold). The tool outputs (i) a text file summarizing the detected blobs and (ii) a PDB file containing the placed water molecules. This tool facilitates the incorporation of structurally meaningful or conserved water molecules into protein–membrane systems, improving model realism for MD refinement within a reproducible Galaxy workflow.
e. **Merge Water Residues tool** extracts selected water residues from a source PDB file and incorporates them into a target system PDB structure. The tool accepts a source PDB containing the residues of interest, a target system PDB, a user-defined list of residue numbers, and a residue-type selector. Selected water residues are appended to the target structure while preserving the original atom and residue numbering. During processing, water residues are converted to the AMBER-compatible WAT naming convention. The output is a single PDB containing the original molecular system together with the user-selected structural waters or other residues, enabling retention of experimentally resolved or otherwise selected waters during subsequent molecular dynamics system preparation.

#### 2.2.3 Galaxy’s Workflow Engine

**GalaxyMemMD** uses Galaxy’s workflow engine, a mature and scalable framework for reproducible biomedical analysis. Galaxy represents workflows as directed acyclic graphs, where each tool is a modular node with explicit inputs, outputs, and versioned metadata. This structure makes each step traceable and allows analyses to be replicated, shared, and inspected with full provenance, which aligns naturally with FAIR principles, meaning data and tools are Findable, Accessible, Interoperable, and Reusable.

A major advantage of the Galaxy implementation is that every stage of protein-membrane modeling is automatically recorded in the Galaxy history. Input datasets, parameter choices, tool versions, intermediate files, and outputs are captured in one place, making the workflow easy to audit, rerun, and modify without ambiguity. Galaxy also supports workflow versioning, so complete pipelines can be stored, published, and reused consistently across projects, research groups, and computing environments.

To further support reproducibility beyond any single machine, the workflow is compatible with containerized execution through Docker and Singularity. This allows each tool to run in a standardized software environment, reducing differences between local systems, cloud platforms, and HPC resources. Combined with provenance tracking, version-controlled workflows, and FAIR-compliant dissemination, Galaxy provides a transparent and shareable framework that is more reproducible and easier to manage than traditional script-based or web-only approaches.

### 2.3 Applying the Workflow

The suite of tools that have been added, and the workflows created with these and Galaxy’s other existing computational chemistry tools, encompass nearly the entire pipeline of steps required for protein-membrane modeling and analysis. For the tested systems, a series of specific steps were used and parameters carefully selected. The primary workflow is divided into 5 subsections, as highlighted in Figure 2. A brief overview of each of the components of implemented workflow are as follows:

**Figure 2.**
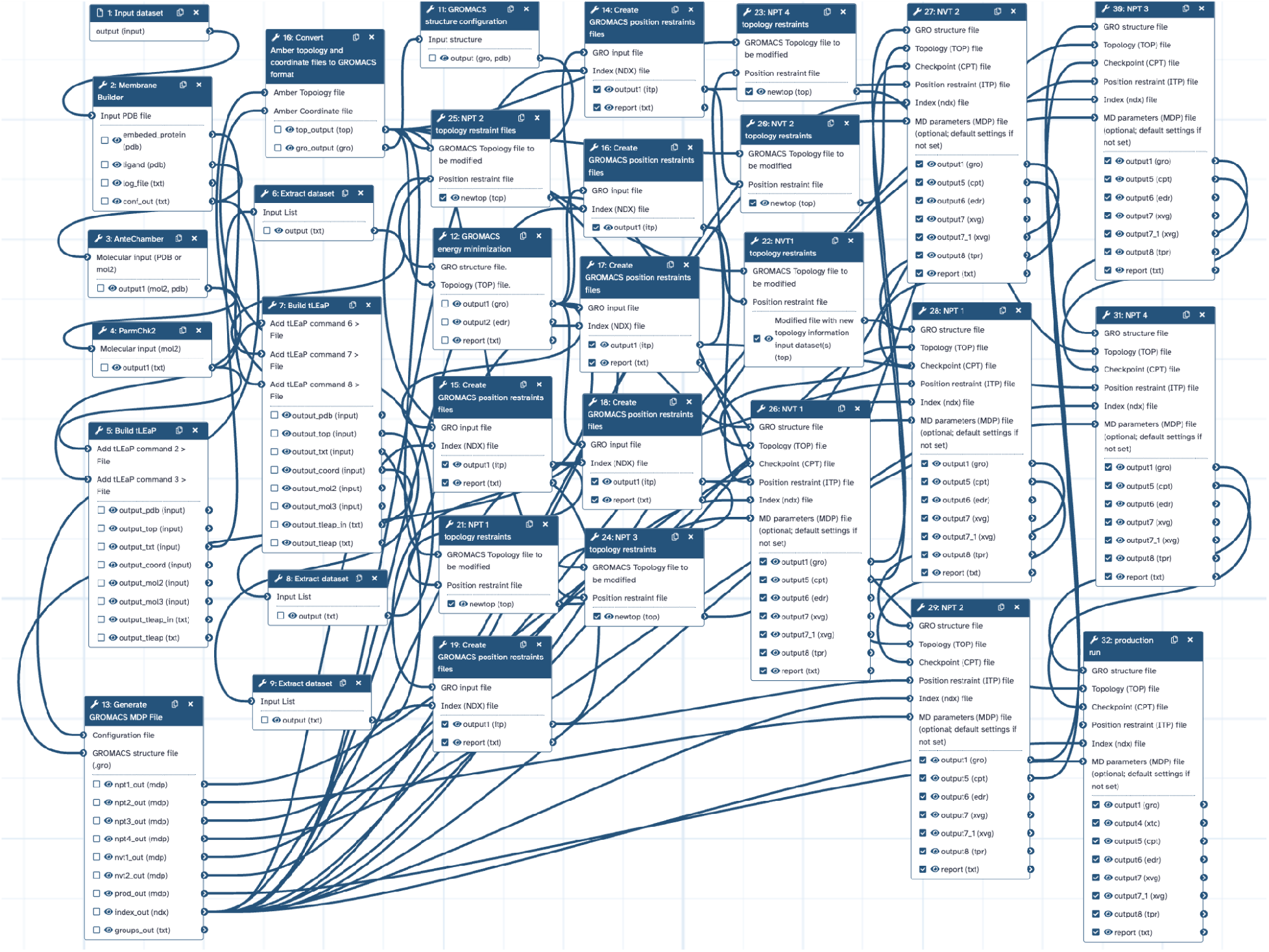
Full node-level view of the Galaxy workflow for membrane protein simulation. Nodes represent Galaxy tools, and edges represent the flow of datasets between them. The pipeline proceeds from membrane embedding and protein/ligand parameterization through energy minimization and equilibration to the production run. All parameters, tool versions, and intermediate datasets are preserved in the Galaxy history, ensuring complete reproducibility.

#### 2.3.1 Membrane Protein Simulation workflow

a. **Membrane Embedding** Membrane embedding step involved Galaxy’s *Membrane Builder tool* to embed both protein–ligand complexes and protein-only systems within a lipid bilayer. Details of membrane composition and dimensions of each simulated system are represented in Table 3. During membrane embedding, the ligand was retained when present, and the tool generated a complete system PDB file, configuration file and a log file documenting the embedding procedure and ligand file. .
b. **Parameterization** Following membrane embedding, the systems were parameterized using Galaxy’s Build tLeap tool ^31^. The protein was described using the ff14SB force field, and TIP3P water was used to solvate the system ^38^. Ligands were parameterized with the second-generation General AMBER Force Field (GAFF2) ^39^, which provides broad coverage for small organic molecules. Ligand partial charges were assigned using the AM1-BCC charge model ^40^ through Antechamber ^41^. Lipid molecules were parameterized using the Lipid17 force field ^42–44^. Together, these force fields and charge models provided a consistent basis for simulating protein–ligand–membrane interactions within the AMBER framework. These settings were used for the benchmark simulations described here; however, the GalaxyMemMD framework allows users to customize their workflows as required.
c. **AMBER-to-GROMACS Trajectory Conversion** Consistent with the framework’s use of fully open-source simulation software, GROMACS ^10^ was used to execute the molecular dynamics simulations after system construction and parameterization with AmberTools. The AMBER topology and coordinate files generated during parameterization were converted into GROMACS-compatible formats using a Galaxy tool that wraps ACPYPE ^45^. ACPYPE enables conversion among AMBER, CHARMM, and GROMACS file formats while retaining the system parameters required for subsequent simulation. A more fine grain conversion is offered by *AMBER to GROMACS tool* where it provides more control over individual system components such as lipid, solvent, ligand and protein by separating out all the components as separate entries in the topology file. The resulting GROMACS structure and topology files were then used to configure the simulation system and generate the position-restraint files required for the staged equilibration protocol.
d. **Energy Minimization** Energy minimization was performed using the GROMACS energy-minimization tool in Galaxy, applying steepest-descent minimization to the complete protein–membrane system. The benchmark workflows used a maximum of 50,000 minimization steps and maximum-force convergence criteria ranging from 100 to 1,000 kJ mol⁻¹ nm⁻¹. Although a high upper limit was specified for the number of steps, each minimization terminated when the selected force-tolerance criterion was reached, ensuring that the system was sufficiently relaxed before molecular dynamics. Electrostatic interactions were treated using smooth particles–mesh Ewald electrostatics (SPME) ^46^, and an initial minimization step size of 0.01 nm was used.
e. **Equilibration and Production Run** After energy minimization, the solvent, ions, membrane, and protein environment were stabilized through a six-stage equilibration protocol. As shown in the workflow, equilibration consisted of two initial simulations in the isothermal–isochoric (NVT) ensemble followed by four sequential simulations in the isothermal–isobaric (NPT) ensemble. Both ensembles employed a stochastic-dynamics integrator, covalent hydrogen-bond constraints using the LINCS algorithm ^47^, and SPME electrostatics. Positional restraints on the protein and lipids were progressively reduced across the six equilibration stages using force constants of 2,000, 1,000, 500, 200, 100, and 50 kJ mol⁻¹ nm⁻². This staged reduction allowed the surrounding membrane and solvent to relax gradually toward an unrestrained state and is consistent with established practices for membrane-protein simulations, including equilibration protocols generated by CHARMM-GUI. Production simulations were initiated directly from the final NPT equilibration stage. All positional restraints were removed, while covalent hydrogen-bond constraints were retained through LINCS. In our benchmark, each production simulation comprised 500 million integration steps with a 2 fs time step, corresponding to a total simulation time of 1 μs per system.
f. **Post-processing and Analysis** Following the production simulations, trajectory files were concatenated, compressed, and processed using *Modify/convert GROMACS trajectories using trjconv and trjcat tool*. The processed trajectories were saved in compressed XTC format and contained 5,000 frames spanning 1 μs for each system. To reduce artifacts associated with periodic boundary conditions, each system was centered on the protein and membrane. These post-processing operations were performed using an additional Workflow, with defined settings for periodic-boundary treatment, unit-cell representation, and centering. The resulting XTC files were suitable for visualization and downstream analysis. Trajectory analysis was performed using another Workflow. The workflow calculated whole-protein root-mean-square deviation (RMSD), whole-protein and site-specific root-mean-square fluctuation (RMSF), radius of gyration (Rg), hydrogen bonds between the ligand and protein-binding-site residues. Ligand-focused analyses included the center-of-mass distance between the ligand and binding-site residues, ligand RMSD, ligand Rg, and dihedral-angle analysis. Galaxy generated standard plots for many of these measurements, while custom Python scripts were used to produce the publication-quality figures presented in the manuscript.

#### 2.3.2 Membrane-embedding Benchmarking

To assess whether each system’s membrane embedding remained physically stable and correctly configured over the ∼1 µs production trajectory, we benchmarked six structural and dynamic parameters using established, published analysis tools. Bilayer integrity was assessed from leaflet composition and lipid packing (leaflet assignment via MDAnalysis LeafletFinder; bilayer thickness via LiPyphilic MembThickness; protein-aware area per lipid via FATSLiM)^48–49^ and acyl-chain order (all-atom S_CD order parameters from explicit C–H bond vectors), reporting on whether the bilayer stayed intact and properly packed rather than thinning, expanding, or disordering. Protein embedding geometry was evaluated from transmembrane-segment identification (DSSP) combined with helix or strand axis tilt relative to the membrane normal (MDAnalysis HELANAL) and insertion depth relative to the bilayer midplane, assessing whether the protein remained correctly and stably inserted rather than tilting away or extruding from the membrane. For the two channel/porin systems (KcsA, OmpG), pore geometry was profiled with HOLE to separate physiological pore/lumen hydration from lipid-matrix water, distinguishing expected channel wetting from a genuine membrane defect. Lipid dynamics were assessed via persistence-filtered flip-flop detection (LiPyphilic) and protein–lipid interface packing via annular contact counts. Each metric was computed as a full-trajectory time series and summarized as block-averaged means to evaluate stationarity, that is, whether the embedded system reached and maintained a stable configuration rather than drifting over the course of the simulation. Details about the method, tool and relevant citation is included in the Supplementary Data S1.

**Table 3.**
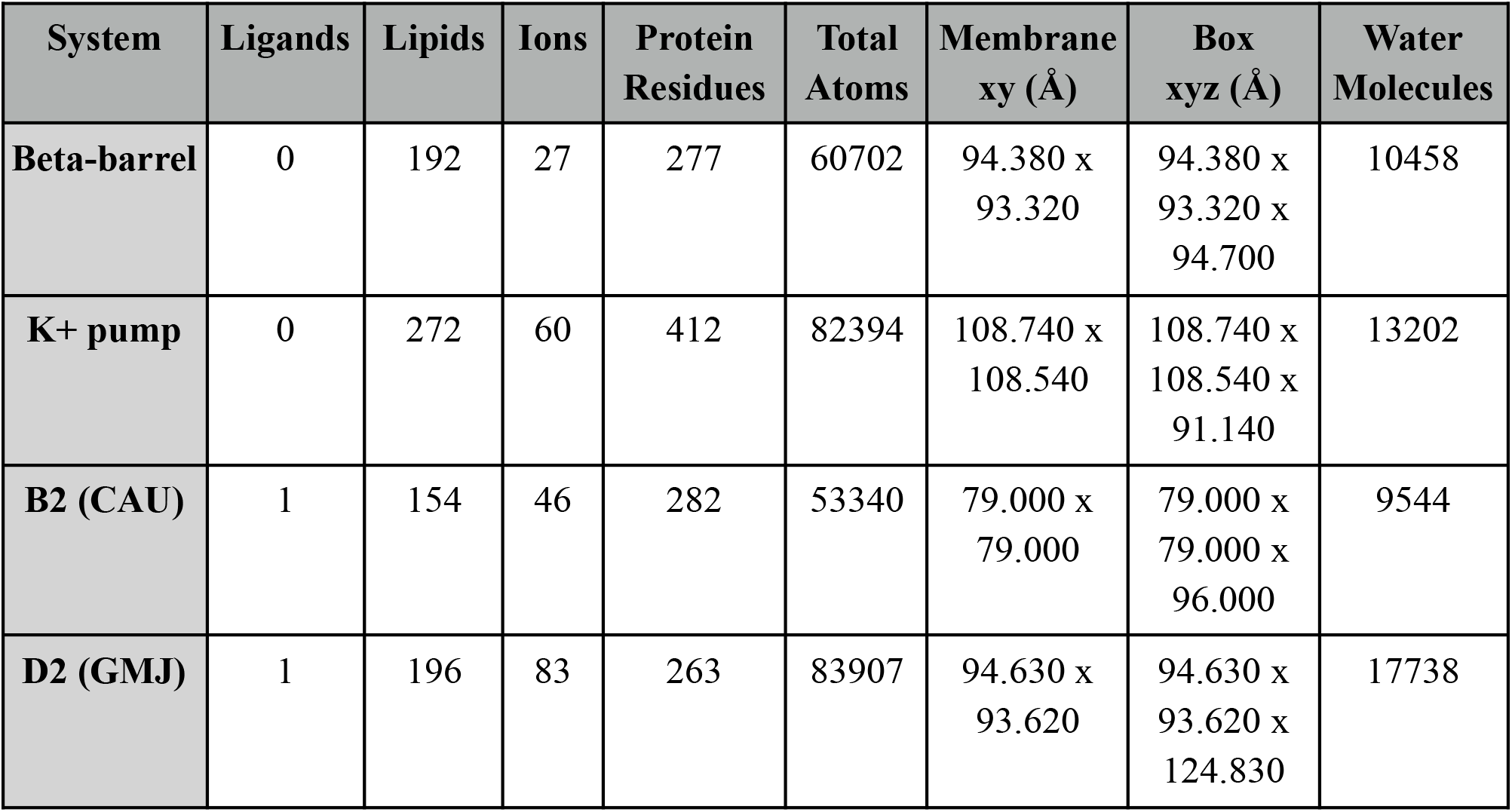
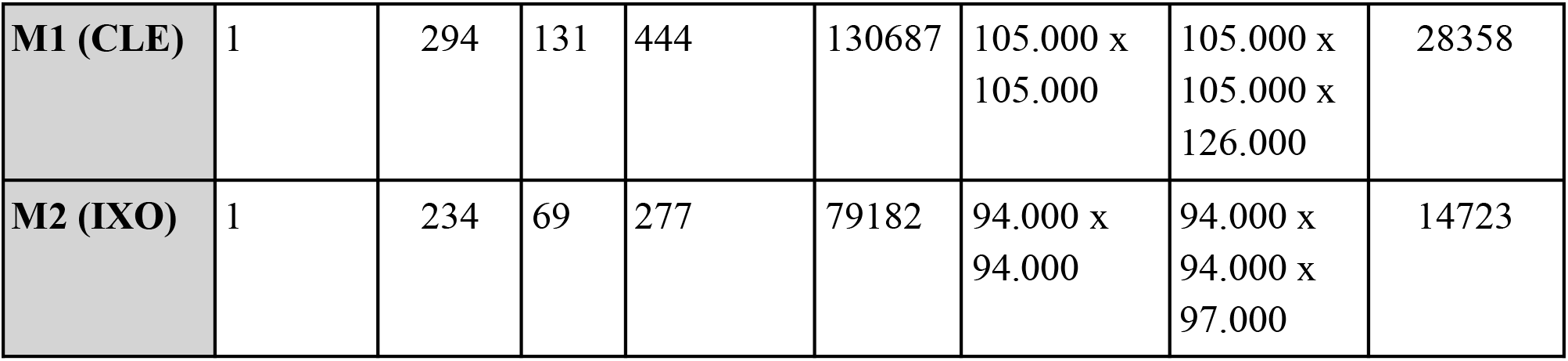
Comparison of the components and dimensions of each simulated system.

| System | Ligands | Lipids | Ions | Protein Residues | Total Atoms | Membrane xy (Å) | Box xyz (Å) | Water Molecules |
| --- | --- | --- | --- | --- | --- | --- | --- | --- |
| Beta-barrel | 0 | 192 | 27 | 277 | 60702 | 94.380 x<br>93.320 | 94.380 x<br>93.320 x<br>94.700 | 10458 |
| K<sup>+</sup> pump | 0 | 272 | 60 | 412 | 82394 | 108.740 x<br>108.540 | 108.740 x<br>108.540 x<br>91.140 | 13202 |
| B2 (CAU) | 1 | 154 | 46 | 282 | 53340 | 79.000 x<br>79.000 | 79.000 x<br>79.000 x<br>96.000 | 9544 |
| D2 (GMJ) | 1 | 196 | 83 | 263 | 83907 | 94.630 x<br>93.620 | 94.630 x<br>93.620 x<br>124.830 | 17738 |
| **M1 (CLE)** | 1 | 294 | 131 | 444 | 130687 | 105.000 x<br>105.000 | 105.000 x<br>105.000 x<br>126.000 | 28358 |
| **M2 (IXO)** | 1 | 234 | 69 | 277 | 79182 | 94.000 x<br>94.000 | 94.000 x<br>94.000 x<br>97.000 | 14723 |

#### 2.3.3 Workflow for 3D-RISM- and Laplacian-guided Placement of Water Molecules

We demonstrate the use of these tools by applying the 3D-RISM- and Laplacian-guided water-placement workflow to the β2-adrenergic receptor (Figure 3).

a. **Preparation of the Protein–membrane System** Initial protein coordinates were supplied as a PDB file or processed using the *Membrane Builder tool* to embed protein with membrane. The ligand was converted into an AMBER-compatible MOL2 representation using Antechamber, while ParmChk2 was used to identify missing force-field parameters and generate the corresponding parameter-modification file. The embedded-protein PDB, ligand MOL2 file, and ligand parameter file were subsequently passed to the *Prepare AMBER System for solvation tool*. This step defined a calculation region based on the solute van der Waals dimensions and produced an AMBER topology (parm7), coordinate file (rst7), LEaP-generated PDB, and cleaned PDB structure (Figure 3).
b. **Calculation of Solvent Density and Placement of Water Molecules** The resulting parm7 and rst7 files were supplied to the Solvent Density Calculator, together with the RISM configuration file and water-model parameter file. A three-dimensional RISM calculation using the Kovalenko-Hirata (KH) closure generated site-resolved solvent maps for the water oxygen and hydrogen sites (kh_O_0.dx and kh_H1_0.dx), as well as the associated 3D-RISM and calculation-information files. The initial oxygen map was then processed using the Solvent Laplacian Calculator, which calculated its spatial Laplacian and generated a refined oxygen map (kh_O.dx) to enhance the localization of discrete hydration sites. The Place Solvent Molecules tool used the initial KH oxygen map, the Laplacian-refined oxygen map, and the AMBER topology and coordinate files to identify clustered solvent-density regions (“blobs”) and position explicit water molecules at the selected sites. This step produced a list of identified density regions (blobs.txt), a PDB file containing the placed waters (in_wat_pdb), and the combined water-containing molecular structure (out_pdb). Finally, the system was reconstructed using Build TleAP, producing the final AMBER topology, coordinate, PDB, MOL2, LEaP input, and LEaP log files. Selected intermediate and final results were collected using the Extract Dataset step for downstream analysis and reproducibility.

**Figure 3.**
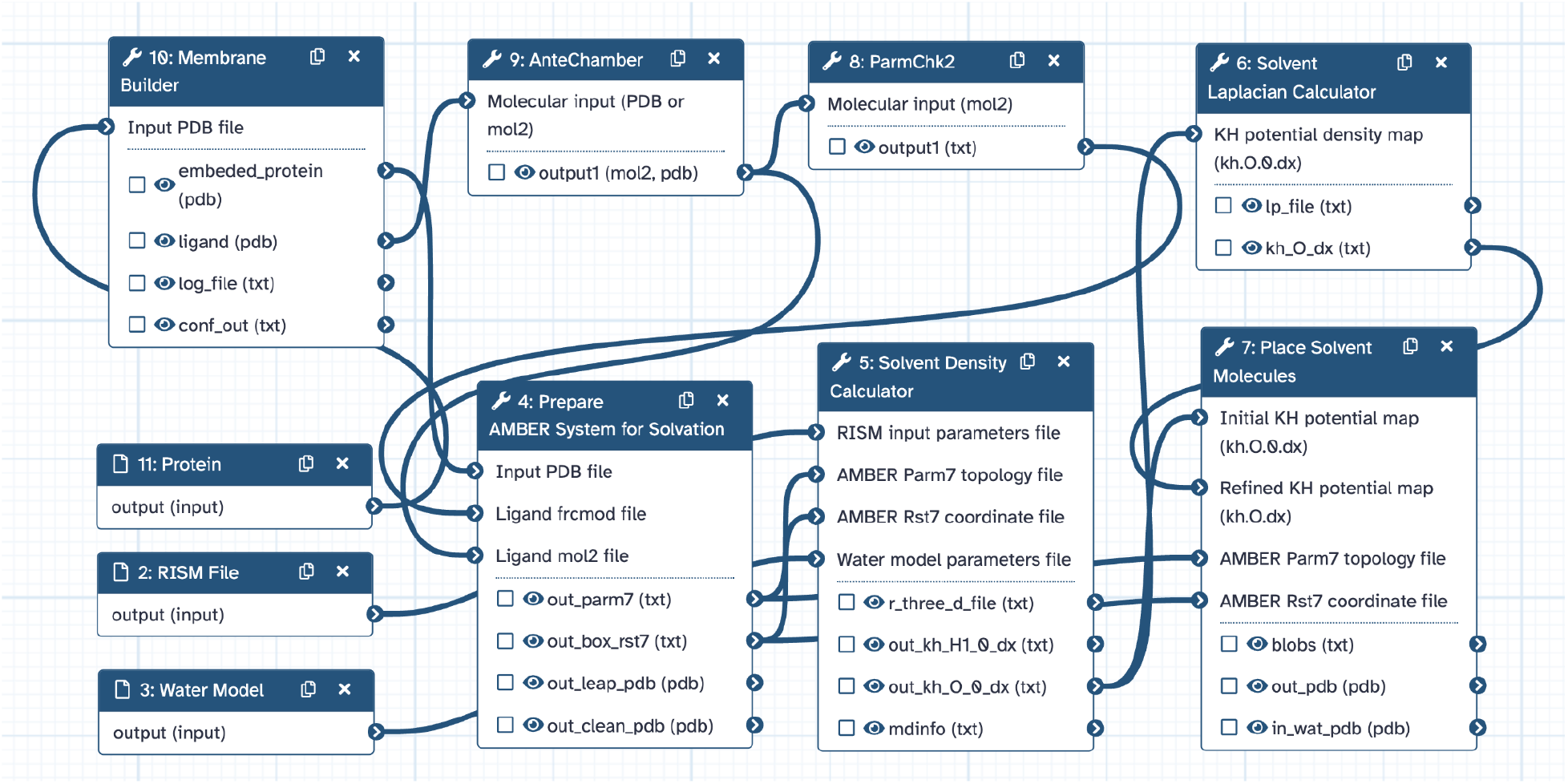
Full node-level view of workflow for 3D-RISM- and Laplacian-guided placement of water molecules. The pipeline proceeds from embedding the protein in the membrane and parameterizing the system, through solvent-density calculation, Laplacian analysis of the density, and hydration-site identification, to the placement of explicit water molecules.

#### 2.3.4 Galaxy Job Execution and Handling

Molecular dynamics simulation is computationally intensive, making efficient resource utilization a critical design consideration. GalaxyMemMD leverages Galaxy’s job execution layer, which is designed for heterogeneous computing environments and enables seamless task submission to local schedulers, HPC clusters, or cloud infrastructures via middleware including SLURM, Pulsar, and Kubernetes. When using GalaxyMemMD on existing public servers (e.g., usegalaxy.org, usegalaxy.eu), users do not need to manage job-handling complexity, as this is maintained by system administrators. For advanced users, Galaxy natively supports a wide range of distributed resource managers (DRMs), including SLURM, PBS/TORQUE, Grid Engine, LSF, and HTCondor, allowing fine-grained configuration via Galaxy’s job configuration system (job_conf.yml / job_conf.xml), where “destinations” define compute resources such as memory, CPU cores, wall-time limits, and queue or partition names. This flexible and modular job management architecture enables parallel execution of multi-membrane simulations, high-throughput workflow replication, and integration with downstream analysis without the knowledge of the complex job execution management systems.

## 3. Results

We tested all three workflows with six different systems. These include four class A GPCR complexes (D2, M1, M2, and B2) and two non-complex proteins (a beta barrel and a potassium pump ion channel). The first workflow involved the creation of the protein-membrane system and subsequent simulation as described above. The second workflow is a GROMACS post-processing workflow to best prepare the trajectories for visualization and analysis. The final workflow is entirely an analysis pipeline developed to measure common metrics used in these types of simulations.

As seen in Table 3, to test customization capabilities of the membrane builder, the simulated systems were constructed with a diversity of dimensions and varying number of components. They range from the smallest case, the B2 complex, which had a total size of 53,340 atoms, to the largest system M1, with 130,687 atoms. The membrane builder allows users to select membrane and box dimensions, which then set the volume for the number of packed lipids and water molecules. Counterions can be added for charge neutralization and additional ions can be included to reflect a specific salinity (default: 0.15 M NaCl). Accordingly, the number of ions in each system ranged from 27 to 131. The GPCRs ranged from 263 to 444 residues in length, with the largest being M1 due to the endolysin fragment included in place of the intracellular domain (ICD). The beta barrel and potassium pump were 277 and 412 residues in length, respectively.

Figure 4a-b summarize the protein-wide stability metrics for the six systems. The C-alpha RMSD traces show distinct stability profiles across the six simulated systems. K+ pump and B2 remain among the lowest-RMSD systems, with mean RMSD values of 1.50 Å and 1.60 Å, respectively. M2 is similarly moderate at 1.80 Å. D2, M1, and beta-barrel show larger average deviations, with mean RMSDs of 2.55 Å, 2.79 Å, and 2.93 Å, respectively. In the case of M1, the structure appears to periodically shift between higher and lower RMSDs, perhaps indicating an oscillation between predominant conformational states. These trends suggest that the final trajectories sample different degrees of global conformational displacement, with the receptor complexes and non-GPCR systems not behaving uniformly under this workflow. The in-depth membrane-embedding benchmarking revealed that all six systems maintained stable bilayer structure and consistent protein-embedding geometry throughout the 1 µs production trajectories, with no evidence of progressive structural drift, membrane defect formation, or protein extrusion (Table S1). A detailed, in-depth benchmarking report is presented in the [Supplementary Data](https://docs.google.com/document/d/1nLBQT8b2lZLTozdISTfOpuEDBbC1fgQKOvN1pyxhOTg/edit?tab=t.0) S1.

**Figure 4.**
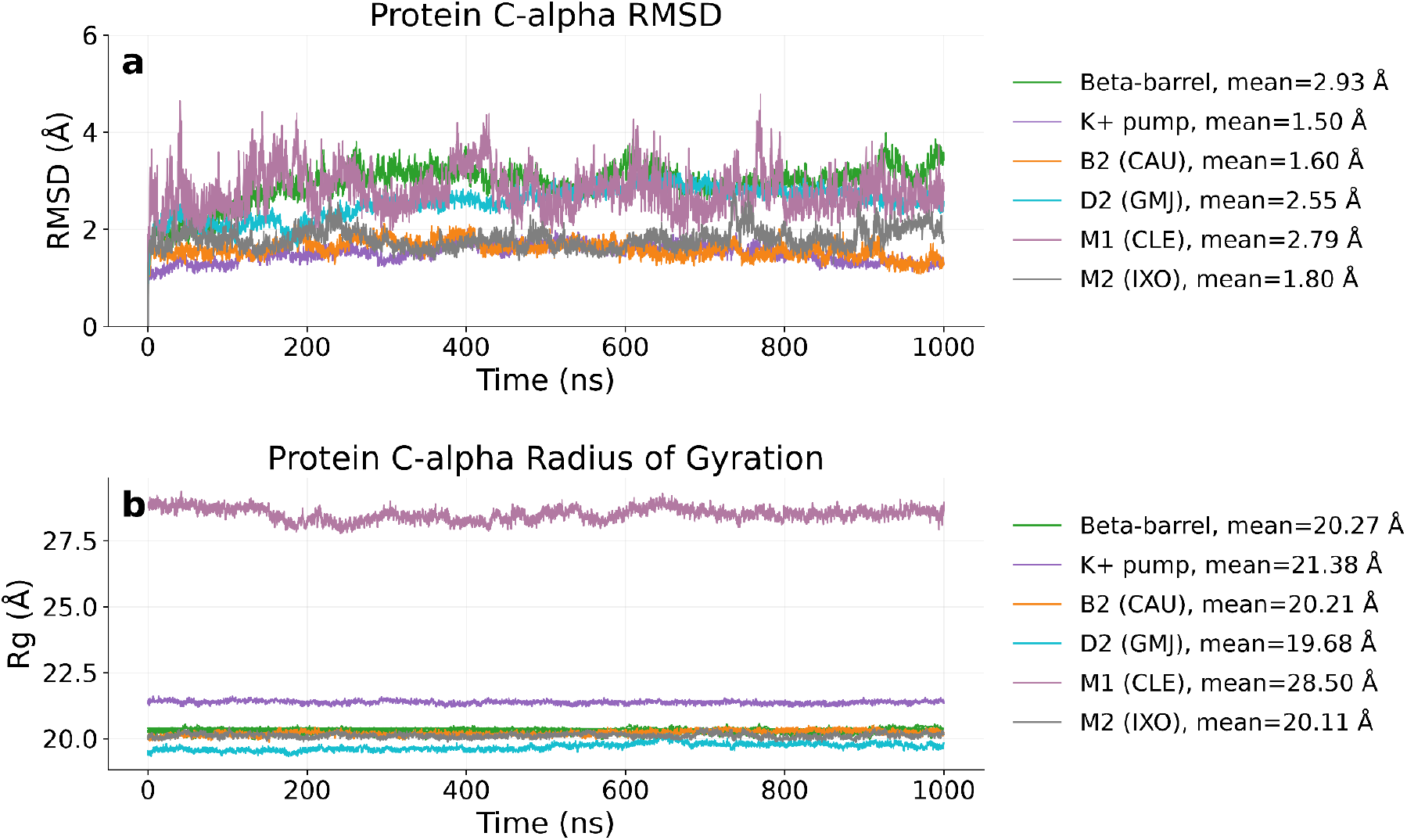
Protein C-alpha structural stability and compactness across the trajectory set. (a) C-alpha RMSD time series for beta-barrel, K+ pump, B2-CAU, D2-GMJ, M1-CLE, and M2-IXO. Mean RMSD values are reported in the legend. (b) C-alpha radius of gyration time series for the same systems. M1 shows a larger whole-construct Rg because the simulated construct contains an endolysin replacement fragment in place of the intracellular domain (ICD), whereas the other receptor systems are truncated. RMSD and Rg are reported in Å.

The protein C-alpha Rg traces show that B2, D2, M2, beta-barrel, and K+ pump remain clustered around approximately 20-21 Å. M1 is a clear outlier, with a mean C-alpha Rg of 28.50 Å. This larger value is expected for the simulated M1 construct because it includes an endolysin replacement fragment in place of the native intracellular domain, whereas the other receptor constructs are truncated. Thus, the M1 Rg should be interpreted as a whole-construct measure rather than a direct comparison of compact receptor-core dimensions.

Figure 5a-f shows the per-residue RMSF profiles for each simulated system. Whole-protein C-alpha RMSF profiles highlight system-specific flexible regions. Mean RMSF values are lowest for K+ pump (0.85 Å), followed by B2 (0.99 Å), M2 (1.02 Å), D2 (1.13 Å), beta-barrel (1.27 Å), and M1 (1.56 Å). The largest local fluctuations are observed in beta-barrel and M2, each exceeding 6 Å at peak residues, while M1 also contains high-fluctuation regions reaching approximately 5.20 Å. These peaks likely correspond to flexible terminal, loop, or construct-specific regions rather than uniform motion across the protein core.

**Figure 5.**
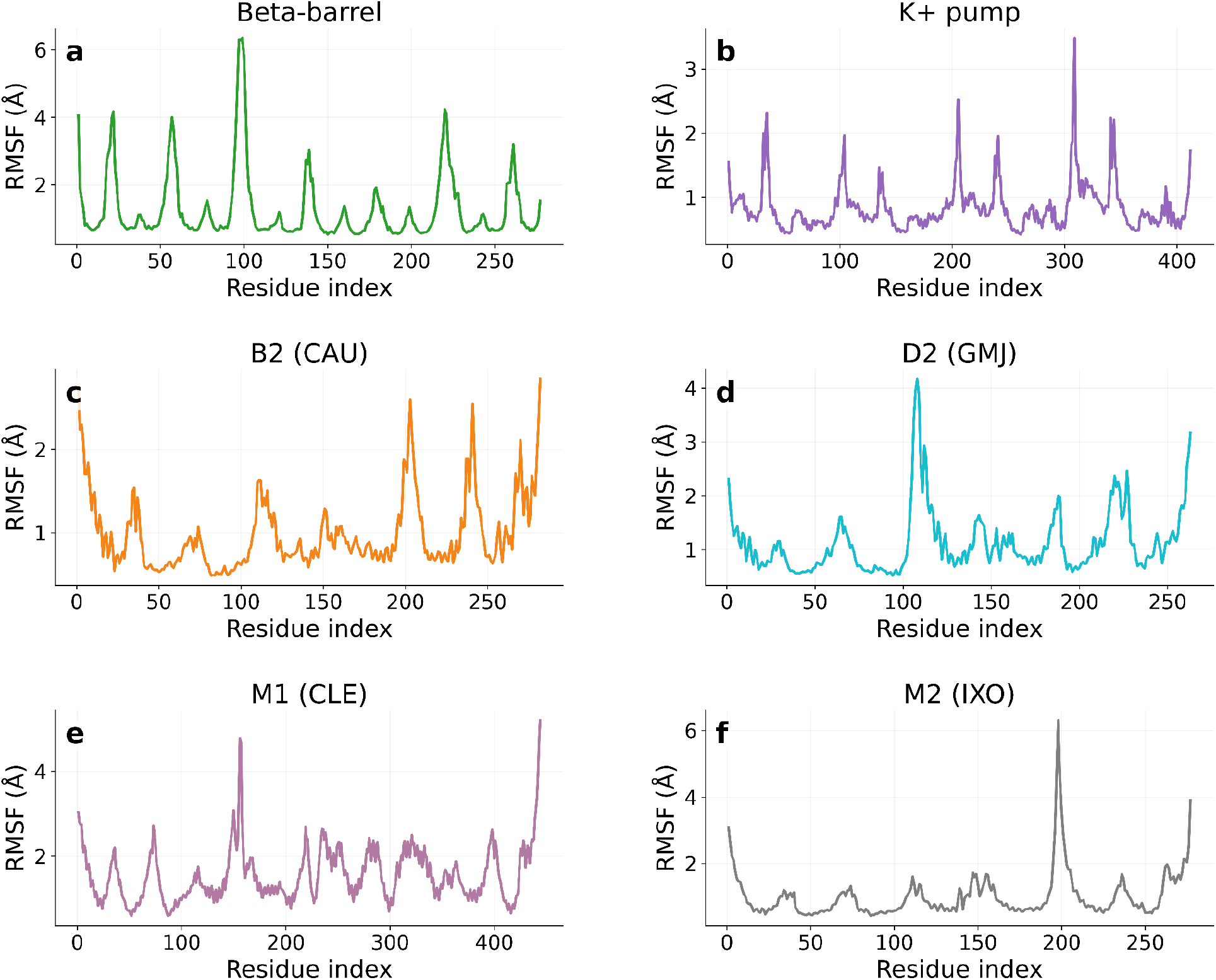
Per-residue C-alpha RMSF profiles for each simulated protein system. Panel images show whole-protein C-alpha RMSF for (a) beta-barrel, (b) K+ pump, (c) B2-CAU, (d) D2-GMJ, (e) M1-CLE, and (f) M2-IXO. RMSF is reported in Å. Residue labels are shown as trajectory residue indices, and system-specific colors are preserved across figures.

The RMSF panel supports using per-system residue-level plots rather than overlaying all systems, because the proteins have different sequence lengths, residue indexing, and construct definitions. The panel format keeps the same visual language and system colors while avoiding direct residue-by-residue comparisons across unrelated indices.

Figure 6a-c summarize ligand position and conformational stability across the four ligand-bound systems. The ligand-to-orthosteric-site COM distance separates the four ligand-bound systems into two broad groups. M1-CLE and M2-IXO remain closer to the orthosteric residue set, with mean COM distances of 2.43 Å and 2.38 Å, respectively. B2-CAU and D2-GMJ show larger mean distances of 4.55 Å and 4.05 Å. D2-GMJ also displays a notable shift late in the trajectory, indicating a change in ligand position relative to the selected orthosteric residues.

**Figure 6.**
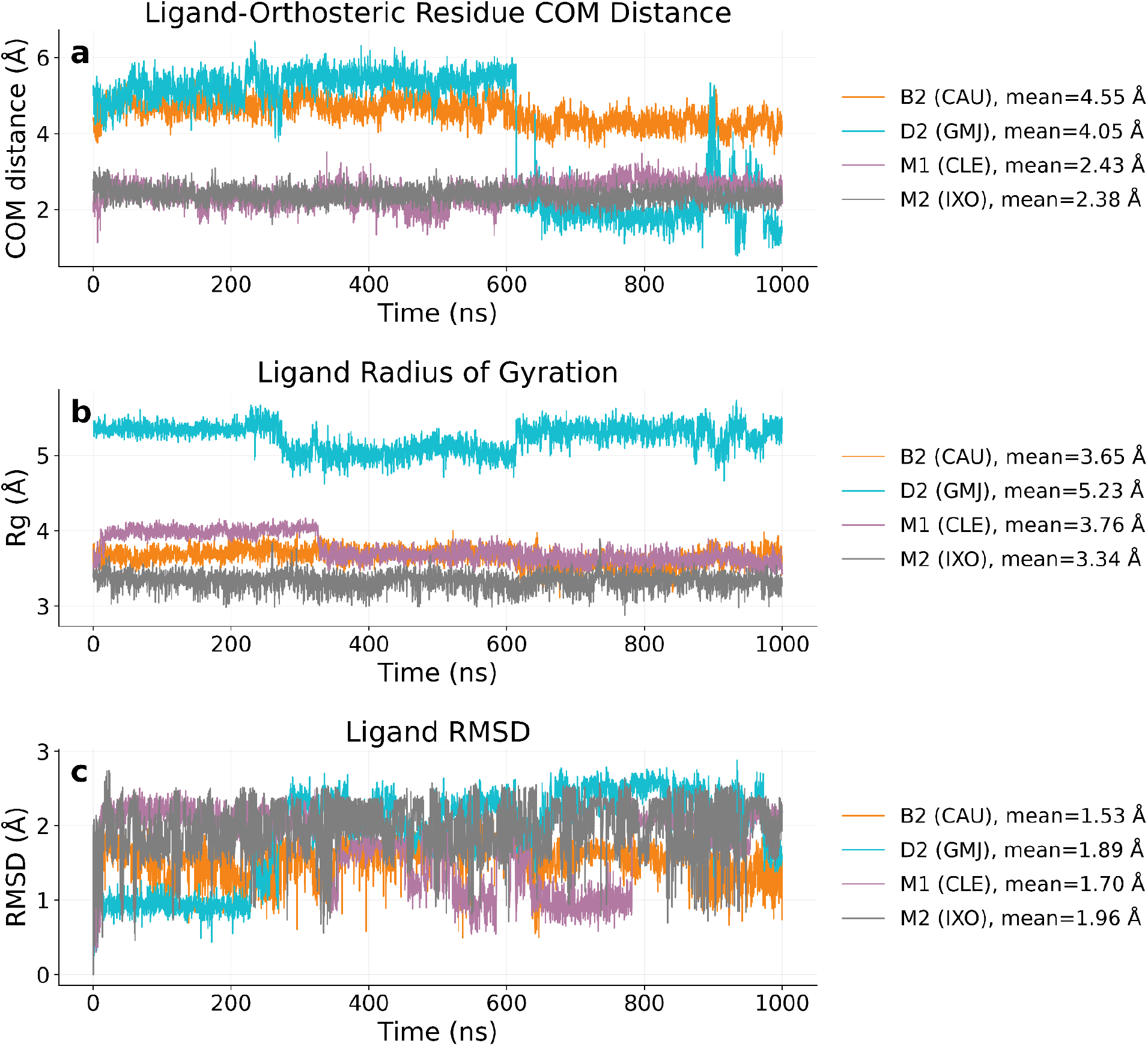
Ligand positional and conformational behavior in the four ligand-bound systems. (a) Center-of-mass distance between each ligand and its corresponding orthosteric residue set. (b) Ligand radius of gyration Rg. (c) Ligand RMSD. Mean values are reported in each legend. CAU is orange, GMJ is cyan, CLE is mauve, and IXO is gray. Distances are reported in Å.

Ligand Rg values suggest that GMJ has the largest and/or most extended ligand geometry in this set, with a mean Rg of 5.23 Å, and a slight transient shift occurring between roughly 300 and 600 nanoseconds. CAU, CLE, and IXO are more compact, with mean Rg values of 3.65 Å, 3.76 Å, and 3.34 Å, respectively. While CAU and IXO retain a more consistent Rg throughout the trajectory, there is notable change in Rg for CLE around 350 ns, but remains consistent thereafter. Ligand RMSD values range from 1.53 Å for CAU to 1.96 Å for IXO, indicating that all ligands show some positional or conformational mobility but remain within a relatively narrow RMSD range for this comparison. The most notable transitions occur with GMJ around 200 ns and with CLE around 300 and again at 800 ns.

Figure 7a-d summarizes the binding-site residue mobility for each ligand-bound system. Binding-site RMSF plots summarize local residue mobility in the orthosteric and allosteric residue selections for each ligand-bound receptor system. Orthosteric site RMSF values are generally modest across the four receptor systems, with M2-IXO showing the smallest orthosteric fluctuations overall. The allosteric selections show more system-specific variability, particularly for M1-CLE, where Gln177 is the most mobile allosteric residue in the selected group. These localized RMSF profiles are useful for assessing whether binding-site residue sets remain stable or contain flexible residues that may influence (or be influenced by) ligand engagement.

**Figure 7.**
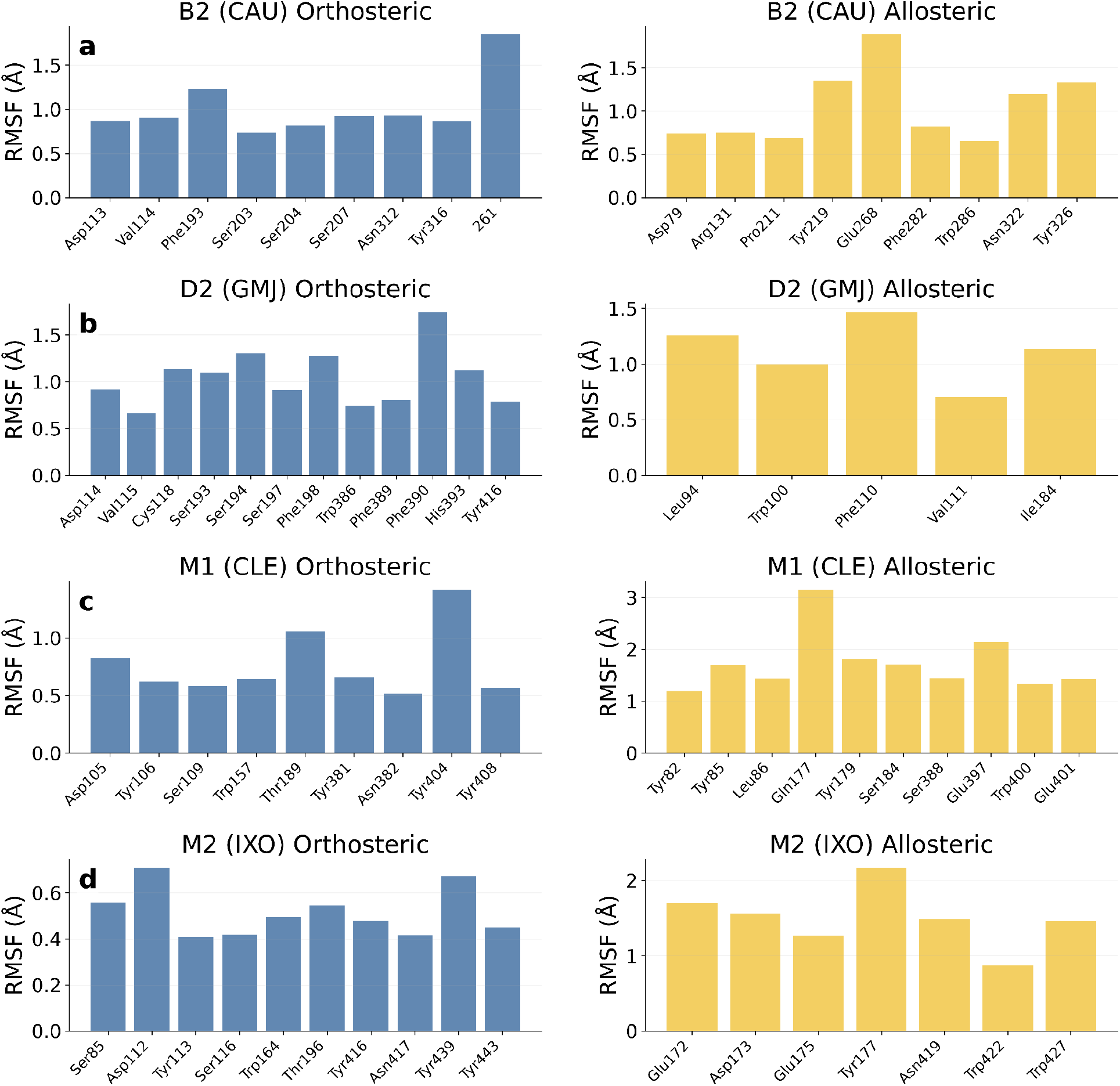
Orthosteric and allosteric binding-site residue RMSF for ligand-bound systems. For each system, orthosteric residue RMSF is shown in blue and allosteric residue RMSF is shown in yellow. Rows correspond to (a) B2-CAU, (b) D2-GMJ, (c) M1-CLE, and (d) M2-IXO. Residue labels use literature/PDB numbering mapped from the simulation .gro residue indices. RMSF is reported in Å.

Residue labels in this panel use the literature/PDB residue names for consistency. Since the workflow’s tools renumber the structure files automatically, which are either truncated to begin with, have an endolysin fragment inserted (M1), or may include missing segments from the N and C termini, a mapping is performed by using dictionaries in a JSON file containing keys and values of the orthosteric and allosteric residue names between both sets of structure files. This makes the binding-site plots more interpretable than the raw .gro residue numbering generated in the workflow, and should be retained for manuscript figures and text.

Figure 8a-d shows the H-bond occupancy rankings and time-series behavior for the ligand-bound systems. Only polar hydrogen bonds are considered in each case, and the cutoff criteria for this analysis was set at a bond angle of 120 degrees and a distance of 3.5 Å. This was done using a VMD-based H-bond analysis Galaxy tool. The analysis shows the strongest and most persistent ligand-protein H-bonding in B2-CAU, with a mean of 5.46 H-bonds per frame and a maximum of 14. D2-GMJ and M2-IXO show intermediate H-bonding, with mean values of 1.67 and 1.31 H-bonds per frame, respectively. M1-CLE has very sparse H-bonding in this trajectory, with a mean of 0.02 H-bonds per frame and a maximum of 1. This is due in part with Clemastine (CLE) only having no hydrogen bond donors and only 2 acceptors, which are largely obscured given the bulky nature of the largely aromatic and cyclic structure.

**Figure 8.**
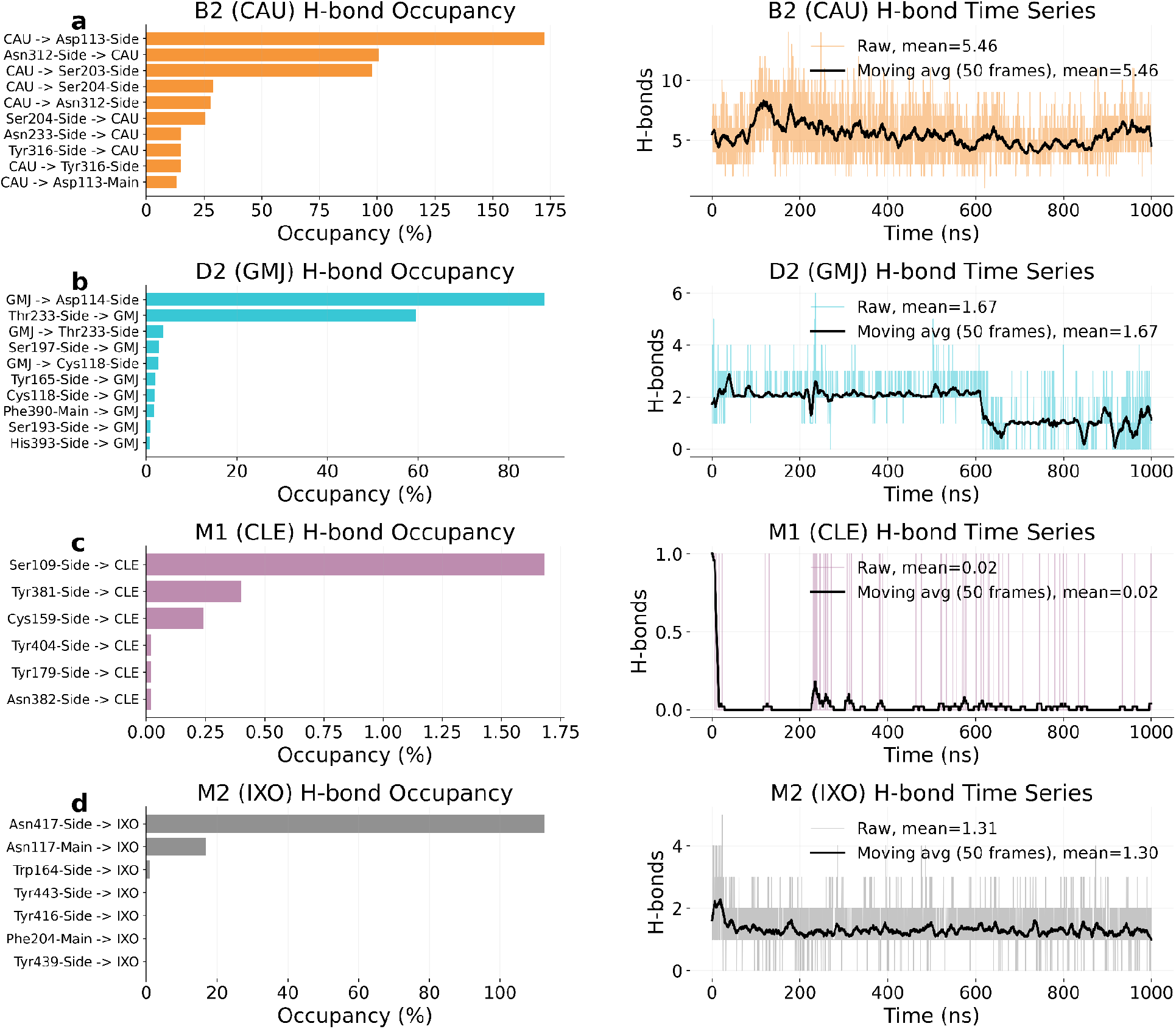
Ligand-protein H-bond occupancy and time-series behavior. For each ligand-bound system, the left panel shows the top H-bond occupancy pairs and the right panel shows the H-bond count time series with a moving-average trace. Rows correspond to (a) B2-CAU, (b) D2-GMJ, (c) M1-CLE, and (d) M2-IXO. Residue labels use literature/PDB numbering where mappings are available. Mean H-bond counts are reported in the time-series legends.

The hydrogen bond occupancy plots show the relative frequency of each donor-acceptor interaction across the trajectory. The left side of the arrow indicates the donor, while the right side is the acceptor. Values that exceed 100% indicate that there could be more than one hydrogen bond contact occurring between the pair. For example, a donor atom on the ligand could be interacting with both a sidechain and backbone acceptor in a nearby residue, or arising from both the ligand and residue pair presenting multiple donor/acceptor atoms each. Occupancy profiles reveal that B2-CAU is dominated by repeated interactions with Asp113, Asn312, Ser203, Ser204, and Tyr316. D2-GMJ is dominated by interactions with Asp114 and Thr233. M1-CLE has low occupancy overall, with Ser109 as the most frequent contact. M2-IXO is dominated by contacts with Asn417 and Asn117. These patterns suggest that the ligand-bound systems differ substantially in polar engagement of the selected binding region, with B2-CAU showing the most extensive H-bond network in the current analysis.

Figure 9a-d shows the ligand images alongside the corresponding dihedral traces. The ligand dihedral traces capture distinct conformational regimes for the four ligands. B2-CAU shows rapid switching across the full angular range, consistent with a highly dynamic torsion that repeatedly crosses the periodic boundary. D2-GMJ remains in a more structured state for much of the trajectory (approximately the first 200 ns and the last 75), with a clear transition from a negative-angle basin to a positive-angle basin and a late trajectory shift to a lower positive range. M1-CLE shows a large consistent rapid fluctuation in dihedral space for the first 320 ns, with a predominantly negative transition between 350 and 550 ns, and the remainder of the simulation presenting a mostly positive angle. M2-IXO is the most broadly distributed of the four traces, with persistent fluctuations across much of the allowed angular space and no single dominant basin. These differences likely arise from both ligand structural diversity and the local binding environment. However, because class A GPCR orthosteric sites are highly conserved, the observed fluctuations are more plausibly attributable to differences in ligand chemistry than to major variation in the binding pocket. The accompanying VMD images provide a useful structural reference for each ligand and make the torsions in the time series easier to interpret. The atom labels in the panel titles document the selected torsion explicitly, and they are colored accordingly in the images.

**Figure 9.**
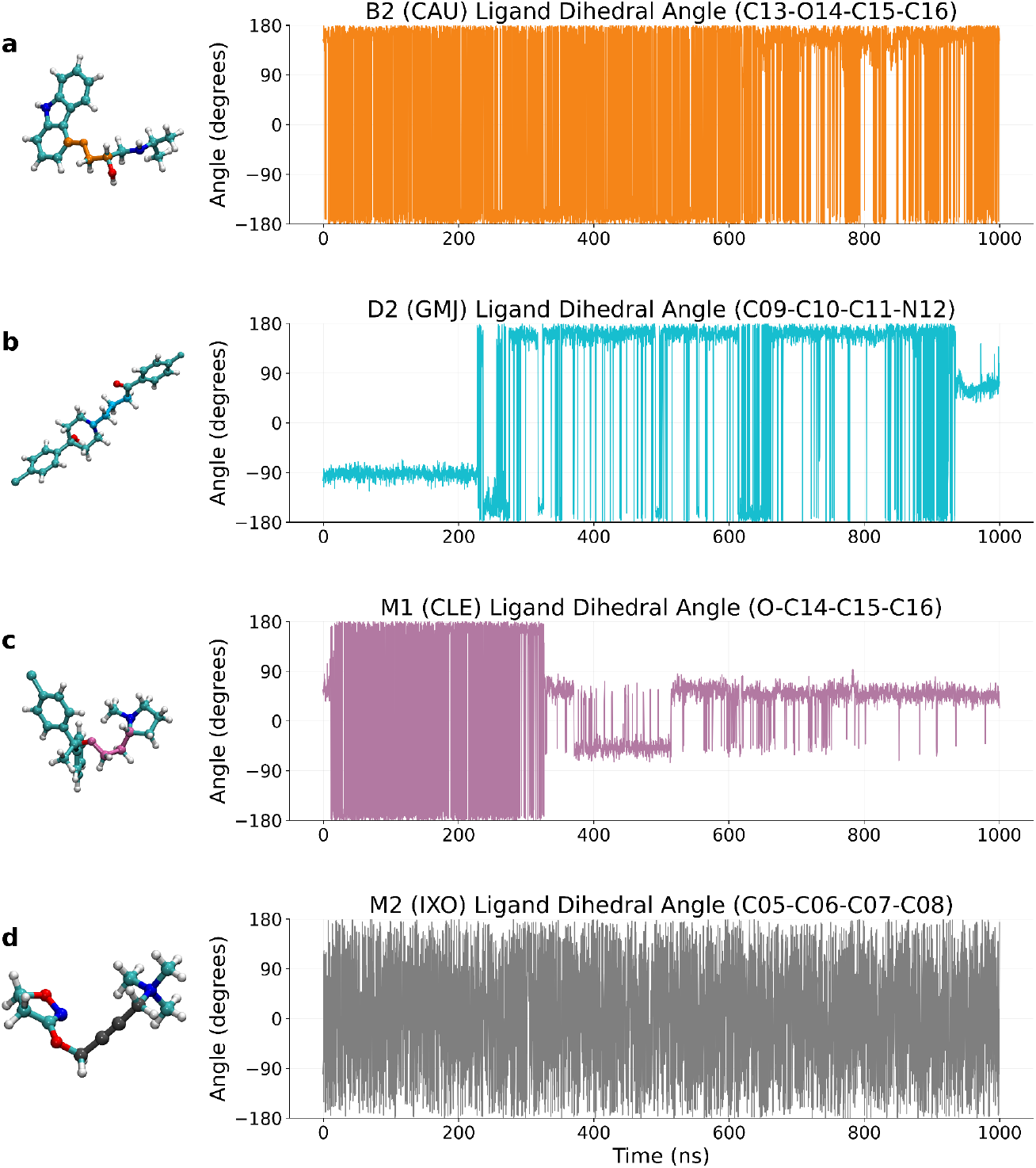
Ligand torsion analysis for the four ligand-bound systems. Each row shows the VMD-rendered ligand image on the left and the corresponding dihedral time series on the right. Rows correspond to (a) B2-CAU, (b) D2-GMJ, (c) M1-CLE, and (d) M2-IXO. The panel titles list the atom names used to define each dihedral angle: B2-CAU, C13-O14-C15-C16; D2-GMJ, C09-C10-C11-N12; M1-CLE, O-C14-C15-C16; and M2-IXO, C05-C06-C07-C08. The M1 trace uses the updated dihedral analysis data from the refreshed archive. Angles are reported in degrees.

In addition, we tested a workflow for predicting solvent distribution around the protein using 3D-RISM calculations. After predicting the positions of water molecules around the protein, we selected individual water molecules based on their location within the protein and their proximity to the ligand (Figure S1), and assessed their behavior during the MD simulations. During the simulation, these two water molecules (resid 16142 and 16143) were tracked throughout the 1 µs MD trajectory using MDAnalysis (Figure S2). Positional stability was evaluated by computing the displacement of each water’s oxygen atom from its initial position at each frame, using a minimum-image-corrected (unwrapped) coordinate scheme to avoid artifacts from periodic boundary crossings. Both water molecules exhibited substantial positional drift over the course of the simulation, with mean displacements of 385.7 Å and 563.6 Å and maximum excursions of 940.7 Å and 1045.3 Å for resid 16142 and 16143, respectively, Figure S2. Neither molecule remained within 5 Å of its starting position for any appreciable fraction of the trajectory (<0.1% of frames), indicating that both waters behaved as freely diffusing bulk solvent molecules rather than as stably bound or structurally ordered waters, consistent with expected bulk-water diffusion over microsecond timescales. Despite their overall mobility, both water molecules made repeated, transient contacts with the protein surface. Heavy-atom contacts (<4.5 Å) with the protein were observed in 11.2% and 10.6% of frames for resid 16142 and 16143, respectively, and hydrogen-bond analysis (donor–acceptor distance <3.5 Å, D–H···A angle >150°) identified 142 and 117 discrete hydrogen-bonding events, corresponding to 2.4% and 2.1% of the simulation time. These transient interactions were not uniformly distributed across the protein surface but were concentrated at a small number of residues, most notably residues 110–113 and 265 for water 16142, and residues 193, 236, 338, and 348 for water 16143, suggesting the presence of preferential, though short-lived, solvent-exposed hydrogen-bonding sites at these positions. In contrast, neither water molecule engaged the ligand CAU to any significant extent: heavy-atom contacts were essentially absent (0.0% of frames), with closest approaches of only 6.16 Å and 2.81 Å, and no hydrogen bonds were formed with the ligand throughout the entire trajectory. Together, these results indicate that the two water molecules sampled here represent mobile bulk-solvent species that transiently interact with the protein surface but do not participate in stable hydration of the CAU binding site.

## 4. Discussion

GalaxyMemMD was developed to eliminate several practical barriers that still constrain protein-membrane simulation workflows. Existing web-based preparation tools are valuable for building starting systems, but they typically stop at setup rather than providing a fully open, end-to-end environment for simulation execution, downstream analysis, and long-term reproducibility. In contrast, GalaxyMemMD is designed to keep the entire workflow inside an open-source, versioned, and shareable Galaxy history, so the full computational record, parameter choices, and outputs remain available after the run rather than being trapped in a transient web session.

The case studies presented here show that this framework is not only more transparent, but also functionally competitive. Across six distinct protein-membrane systems, the workflow produced stable embedded systems, reproducible trajectory outputs, and interpretable structural metrics without requiring manual intervention at each stage. The system overview panel confirms that the final prepared complexes are visually consistent with the intended membrane environments, while the structural analyses in Figures 4 through 9 show expected behavior across a diverse set of membrane proteins and ligands. Protein RMSD and radius of gyration values remain in biologically reasonable ranges for most systems, residue-level RMSF profiles resolve system-specific flexible regions, ligand COM distances and ligand conformational metrics separate the ligand-bound systems in meaningful ways, and the H-bond and dihedral analyses provide additional orthogonal views of binding behavior and ligand dynamics.

A key advantage of GalaxyMemMD is that it supports not just simulation setup, but also the analysis layer that follows it. This is where many existing tools become limiting. Users can prepare a system in a web interface, but then they typically need to export files, move them elsewhere, reconstruct the analysis environment, and manually track parameters and file versions. GalaxyMemMD avoids that fragmentation. The workflow can retain the prepared structure, the simulation inputs, the production trajectories, and the analysis outputs in one place, which is particularly important for long-term storage, review, and reuse in a publication setting. That continuity is not a minor convenience; it materially improves reproducibility and makes the workflow more suitable for collaborative projects.

The comparison figures also show that the workflow is flexible enough to handle systems with different topologies and different analysis needs. The beta-barrel, K+ pump, and GPCR systems do not have the same construct boundaries, residue numbering, or ligand definitions, yet the workflow still produces coherent overlays and system-specific results. Residue renaming through JSON dictionaries, separate orthosteric and allosteric analyses, and per-system ligand annotations all make the output interpretable without requiring custom one-off scripts for each target. That matters for practical use, because protein-membrane projects rarely stay uniform across targets.

We attempted to validate our efforts by comparing our results to previous simulations that were built using CHARM-GUI. In particular for the M1-CLE system, the results indicate that there is overlap between the older simulations in prior work ^50^. Specifically, the new data measures ligand COM distance at 2.23 Å (2.34 Å previously), ligand Rg at 3.76 Å (3.78 Å previously), and whole protein CA Rg at 2.850 Å (2.876 previously). The orthosteric residue RMSFs also demonstrate similar mobility and both the current and previous simulations establish Tyr404 as the dominant flexible residue. There is less agreement in the ligand RMSD, measured at 2.79 Å in the current simulations (1.98 Å previously), and the whole protein CA RMSD measured at 1.70 Å (2.45 Å previously). One important distinction is that the previous data is averaged over three replica simulations, while the work presented here involves data from a single replica simulation per system. Nonetheless, the consistency in some of these key metrics reaffirms the reproducibility in our approach at building, parameterizing, and simulating these systems.

The dihedral analysis is a useful example of why the integrated approach matters. Once the ligand geometries were visualized and the correct torsions were used, the results show both the molecular context and the time evolution side by side. That kind of compact, auditable result is much harder to maintain in a fragmented workflow where plotting, structure rendering, and trajectory analysis live in separate environments. The same is true for the H-bond analysis, where occupancy and time series are presented together and can be compared directly across systems.

Notably, in our second most important workflow in this project is prediction of solvent around protein and the two water molecules analyzed here (resid 16142 and 16143) were selected arbitrarily through visual inspection of the simulation, without any prior knowledge of their proximity to functionally relevant sites or their potential role in ligand binding. This unbiased selection provides a useful stress test of the water-tracking workflow’s ability to correctly characterize hydration behavior in the absence of any expectation bias. The workflow correctly identified both molecules as mobile, bulk-like solvent; their displacement profiles were consistent with diffusive behavior expected for water molecules with no structural role, rather than producing spuriously "stable" trajectories that might arise from artifacts in trajectory unwrapping or coordinate handling. At the same time, the pipeline remained sensitive enough to detect and quantify genuine, geometrically valid transient hydrogen-bonding events with specific protein surface residues, rather than returning a uniformly null result, demonstrating that it can resolve short-lived, biologically plausible interactions even for waters with no a priori significance. Equally important, the workflow did not report spurious contacts or hydrogen bonds with the CAU ligand despite the water molecules sampling a large volume of the simulation box over the 1 µs trajectory, indicating that the distance and geometric criteria applied are appropriately conservative and not prone to false-positive assignment of ligand engagement.

Taken together, these observations support the robustness of the workflow as a general-purpose tool for characterizing water behavior around the protein–ligand system, independent of whether the water molecules under investigation are pre-selected for known biological relevance. This is particularly relevant for downstream application of the pipeline to water molecules identified through predictive or automated water-mapping approaches, where the biological significance of candidate hydration sites is not known in advance.

Taken together, these results support the central claim of the project: a fully open-source Galaxy-based workflow can match the practical utility of more closed or partially closed protein-membrane preparation systems while extending beyond preparation into simulation handling, provenance, and analysis. In other words, the value of GalaxyMemMD is not only that it lowers the technical barrier to entry, but that it preserves the scientific object of the analysis across the full lifecycle of the project. For protein-membrane studies where reproducibility, reanalysis, and long-term accessibility matter, that end-to-end design is the real advantage.

Several limitations should be considered when interpreting this study. The benchmark includes six membrane proteins and a limited set of lipid compositions, with one production trajectory per system; it therefore demonstrates workflow feasibility and provenance rather than statistical convergence or broad biological generalizability. We did not perform a controlled head-to-head assessment of setup accuracy, runtime, resource use, or user effort against other platforms, and the 3D-RISM-guided water-placement procedure was not independently validated against experimental hydration sites. Future work should expand protein and membrane diversity, include independent simulation replicas and convergence analyses, and provide standardized performance and accuracy benchmarks.

## 5. Data availability

The GalaxyMemMD tools and workflows are open-source and available on GitHub under the MIT license at https://github.com/jaidevjoshi83/GalaxyMemMD. The tools can be installed on any Galaxy instance from the Galaxy ToolShed and are available for use on the public server usegalaxy.eu. The Galaxy workflow files (.ga), the starting protein structures, the MD trajectories and topologies, and the analysis outputs needed to reproduce the results presented here are available in Zenodo at https://doi.org/10.5281/zenodo.22967526. The workflows can be imported into any Galaxy instance and run with the provided input structures to reproduce the analyses or to apply the pipeline to new membrane proteins.

## Author Contributions

J.J. conceived the research. J.J. and B.R. designed and implemented the tools and workflows. N.K. performed testing and benchmarking of the tools and workflows. B.R. ran the molecular dynamics simulations and analyzed the data. D.B. supervised the research. D.B. and F.C. provided critical feedback on the study design and manuscript. J.J., B.R., N.K., F.C., and D.B. wrote the manuscript. All authors read and approved the final version.

## Acknowledgments

We would also like to acknowledge the use of AI for language editing to improve clarity and readability, limited to sentence structure and word choice suggestions. The research ideas, data analysis, interpretations, and conclusions presented here are solely the product of the authors’ analysis and expertise.

---

**Figure S1.**
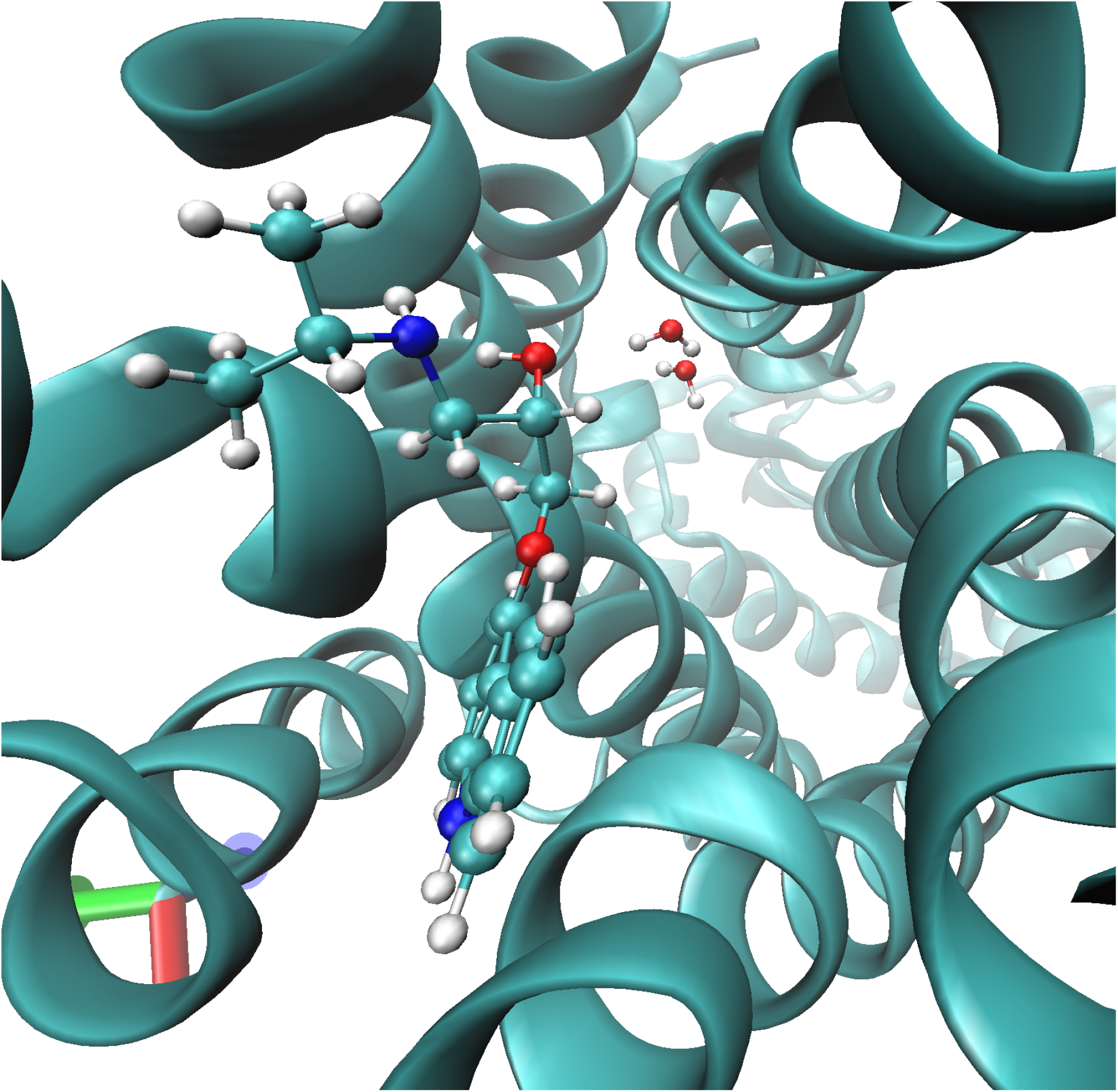
Predicted water molecules within the ligand-binding site. Hydration sites were identified using the 3D-RISM–Laplacian workflow. Two water molecules located near the bound ligand were selected for subsequent assessment of their occupancy and persistence within the binding pocket. The protein is shown as a cyan cartoon, while the ligand and selected water molecules are shown as sticks and ball-and-stick representations, respectively.

**Figure S2.**
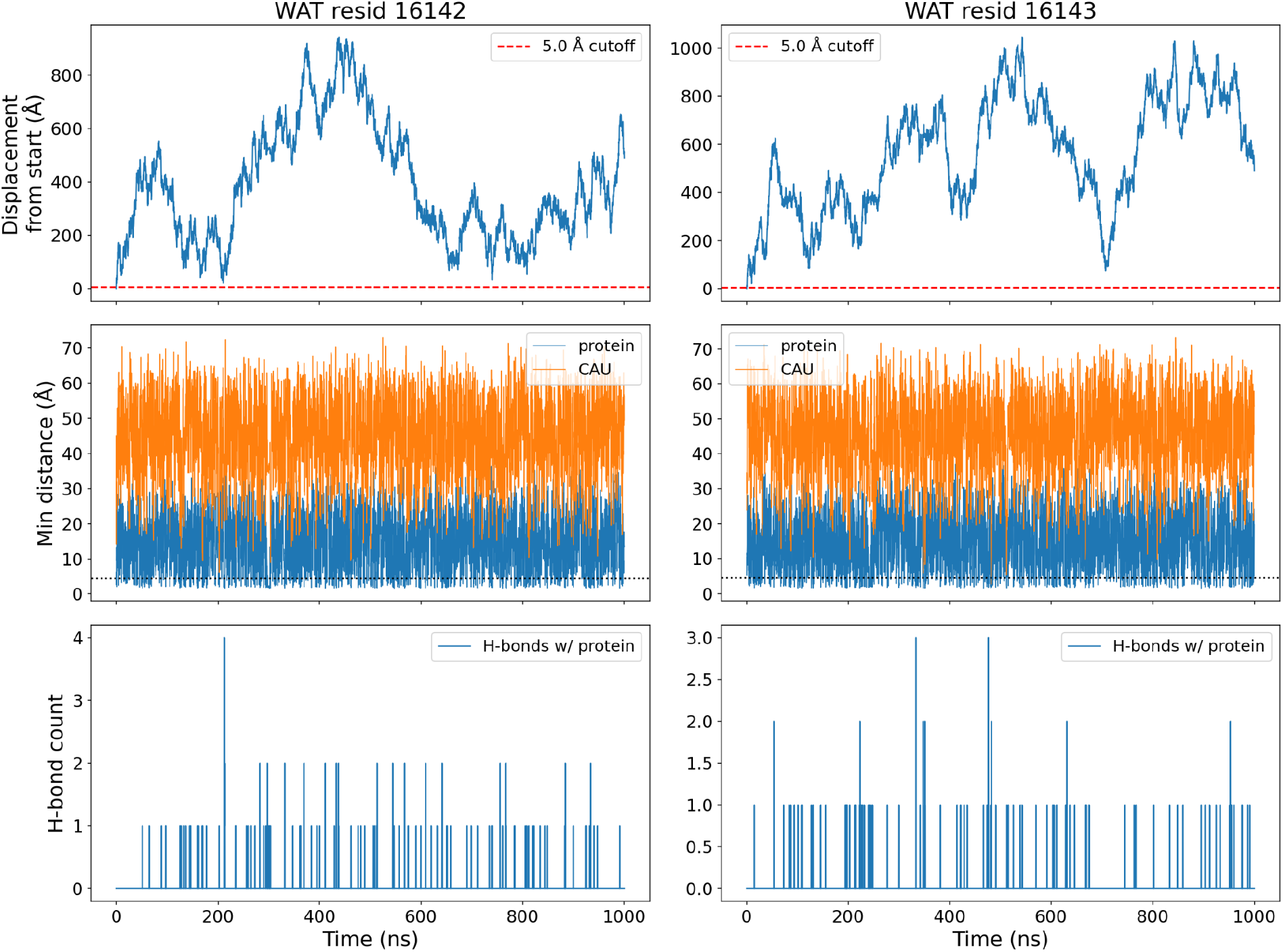
Positional stability and interaction dynamics of two arbitrarily selected water molecules (WAT resid 16142, left; resid 16143, right) over a 1 µs MD trajectory. Top row: Displacement (Å) of each water’s oxygen atom from its initial position, computed from minimum-image-unwrapped coordinates to correct for periodic boundary crossings; the dashed red line marks a 5 Å reference cutoff, below which a water molecule would be considered positionally stable. Middle row: Minimum heavy-atom distance (Å) between each water molecule and the protein (blue) or the ligand CAU (orange) at each frame; the dotted black line marks the 4.5 Å heavy-atom contact cutoff. Bottom row: Number of hydrogen bonds formed per frame between each water molecule and the protein, no hydrogen bonds were detected between either water molecule and CAU over the course of the simulation. Time is shown in nanoseconds (0–1000 ns, corresponding to 5001 trajectory frames saved every 200 ps).

