## Supplementary Data S1 for "GalaxyMemMD: an end-to-end Galaxy toolset for reproducible membrane embedding, molecular dynamics simulation, and analysis of membrane proteins"

**Supplementary Data S1** This supplementary document reports membrane-embedding validation analyses for six GalaxyMemMD production trajectories (B2-CAU, D2-GMJ, KcsA, M1-CLE, M2-IXO, OmpG).

#### Membrane-Embedding Validation

Benchmark analyses presented in this study support the physical consistency and membrane-embedding stability of the six simulated systems. All systems maintained stable membrane-embedding characteristics throughout the simulations, with membrane thickness fluctuating by only 2–5% around the mean (standard deviations of 0.07–0.10 nm for mean thicknesses of 3.70–3.88 nm). Similarly, the projected membrane area exhibited fluctuations of approximately 1–2%, with no evidence of progressive membrane collapse or excessive expansion. The area per lipid (APL) values were also consistent with the expected physicochemical properties of the constituent lipids. Specifically, the two POPC-containing systems (D2-GMJ: 0.71–0.72 $nm^2$; KcsA: 0.69–0.70 $nm^2$) exhibited systematically higher APL values than the four DPPC-containing systems (0.61–0.68 $nm^2$), consistent with the expected differences in lipid packing. FATSLiM-based analysis therefore reproduced the anticipated lipid chemistry-dependent ordering across independently constructed systems.

Protein tilt angles were similarly consistent with the expected structural characteristics of the different membrane-protein folds. The four GPCR systems (β2, D2, M1, and M2) exhibited tilt angles within a relatively narrow range of 20–24°, indicating consistent membrane-embedding geometries among independently constructed seven-transmembrane receptors. In contrast, KcsA and OmpG exhibited larger tilt angles of approximately 35° and 39°, respectively. The higher average tilt observed for KcsA may partly reflect the inclusion of more obliquely oriented pore helices, whereas the larger value observed for OmpG may reflect differences associated with its β-barrel architecture and the corresponding definition of the structural axis. These fold-dependent differences indicate that the analysis captures meaningful structural variation rather than arbitrary fluctuations. All six proteins also remained stably embedded within their respective membranes, with insertion depths remaining within ±0.6 nm of the bilayer midplane, substantially smaller than the approximately 1.8–1.9 nm bilayer half-thickness, and with no evidence of protein extrusion or progressive displacement from the membrane.

Lipid flip-flop analysis further supported membrane stability, with three of the six systems exhibiting no persistent flip-flop events after application of a 5-ns persistence criterion. Finally, core hydration was consistent with the geometrical properties of the membrane channels. The narrow KcsA pore, with a HOLE-derived radius of 0.235 nm, contained a low and stable average water population of approximately 0.9 molecules, whereas the substantially wider OmpG pore (1.715 nm) maintained an average of approximately 88.7 water molecules. Together, these observations demonstrate that the benchmark analyses recover expected relationships among membrane structure, lipid composition, protein architecture, and channel geometry, supporting the physical consistency and membrane-embedding stability of the simulated trajectories. (Figure S3.1-8.4).

| Parameter | B2-CAU | D2-GMJ | KcsA (1R3J) | M1-CLE | M2-IXO | OmpG |
|---|---|---|---|---|---|---|
| **Lipid species** | DPPC | POPC | POPC | DPPC | DPPC | DPPC |
| **N lipids (up/low)** | 154 (77/77) | 196 (99/97) | 272 (132/140) | 294 (147/147) | 234 (118/116) | 192 (94/98) |
| **Thickness (nm)** | 3.88 +/- 0.09 | 3.71 +/- 0.07 | 3.70 +/- 0.07 | 3.80 +/- 0.09 | 3.81 +/- 0.10 | 3.88 +/- 0.08 |
| **Proj. area (nm²)** | 55.38 +/- 1.14 | 74.15 +/- 1.10 | 104.54 +/- 1.38 | 100.51 +/- 1.67 | 81.10 +/- 1.42 | 69.68 +/- 1.54 |
| **FATSLiM area/lipid (nm², up/low)** | 0.634 / 0.620 (up/low) | 0.713 / 0.715 (up/low) | 0.696 / 0.694 (up/low) | 0.628 / 0.654 (up/low) | 0.675 / 0.653 (up/low) | 0.614 / 0.626 (up/low) |
| **HELANAL tilt (deg)** | 23.67 +/- 3.32 | 20.54 +/- 2.68 | 35.15 +/- 0.82 | 20.00 +/- 1.28 | 21.81 +/- 1.98 | 38.68 +/- 1.06 |
| **Insertion depth (nm)** | -0.541 +/- 0.224 | 0.481 +/- 0.164 | -0.059 +/- 0.120 | 0.529 +/- 0.136 | 0.586 +/- 0.118 | -0.542 +/- 0.171 |
| **LiPyphilic flip-flops** | 0 (>=5ns persistence) | 2 (>=5ns persistence) | 0 (>=5ns persistence) | 18 (>=5ns persistence) | 2 (>=5ns persistence) | 0 (>=5ns persistence) |
| **Core water** | 32 (median, non-pore) | 8 (median, non-pore) | pore=0.9, matrix(median)=6.0 | 12 (median, non-pore) | 11 (median, non-pore) | pore=88.7, matrix(median)=4.0 |

***Table S1. Cross-system benchmark of membrane-embedding validation metrics, six GalaxyMemMD production trajectories (1000 ns each).*** *All values are mean +/- standard deviation (structural fluctuation over the trajectory) unless stated otherwise; tools are as listed in the Methods and citations table below.*

**Lipid species -** bulk membrane lipid, verified independently from each system's own bonded topology.

**N lipids (up/low)** - total reconstructed complete lipid molecules, with the upper/lower leaflet split from MDAnalysis LeafletFinder.

**Thickness (nm)** - bilayer thickness, LiPyphilic MembThickness.

**Proj. area (nm2)** - projected membrane area from the simulation box vectors (Lx x Ly)

**FATSLiM area/lipid (nm2, up/low)** - protein-aware area per lipid (2D Voronoi tessellation against the protein footprint), FATSLiM, reported separately for the upper and lower leaflet.

**HELANAL tilt (deg)** - mean acute angle between the transmembrane-domain axis and the instantaneous membrane normal, averaged over all DSSP-identified transmembrane segments per system: MDAnalysis HELANAL axes for alpha-helices, or the direct first-to-last C-alpha vector for beta strands.

**Insertion depth (nm)** - signed projection of the transmembrane-domain C-alpha centroid onto the membrane normal, relative to the bilayer midplane. Sign, +/- , reflects displacement toward one leaflet or the other, not a directional error.

**LiPyphilic flip-flops** - number of genuine leaflet-to-leaflet relocation events over the full trajectory (LiPyphilic FlipFlop, requiring >=5 ns continuous residence in the new leaflet before counting an event, which filters out transient interfacial noise).

**Core water** - water-oxygen occupancy within +/-0.5 nm of the instantaneous bilayer midplane.

#### Method, tools and citations

| Analysis | Tool |
|---|---|
| Trajectory handling / atom selection | MDAnalysis[1] |
| Bilayer thickness | LiPyphilic MembThickness[2] |
| Area per lipid (protein-aware) | FATSLiM[3] |
| Acyl-chain order parameters (S_CD) | Direct C-H bond-vector calculation from bonded topology[4] |
| Transmembrane secondary-structure identification | DSSP (mkdssp)[5,6] |
| Helix axis / tilt | MDAnalysis HELANAL[1,7] |
| Pore radius profiling (KcsA and OmpG) | HOLE (via MDAnalysis hole2 interface)[8] |
| Flip-flop detection | LiPyphilic[2] |
| Protein-lipid contacts (0.5 nm heavy-atom cutoff) | MDAnalysis capped_distance[1,2] |
| Density profiles | Direct histogram along membrane normal[9] |

***Table S2****. Analysis method, tool and references.*

### B2-CAU-membrane System

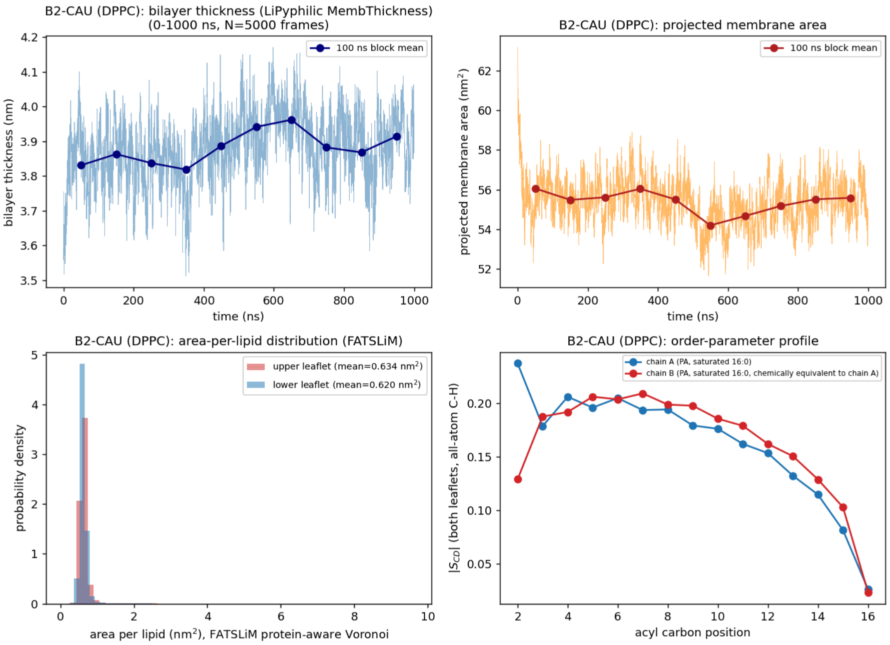


**Figure S3.1.** *Membrane structure. Bilayer thickness and projected membrane area vs. time (raw trace with 100 ns block means); area-per-lipid distribution by leaflet (FATSLiM, protein-aware Voronoi tessellation, Buchoux 2017); mean |S_CD| (absolute value of the deuterium order parameter) acyl-chain order-parameter profile for both lipid tails, from explicit C-H bond vectors.*

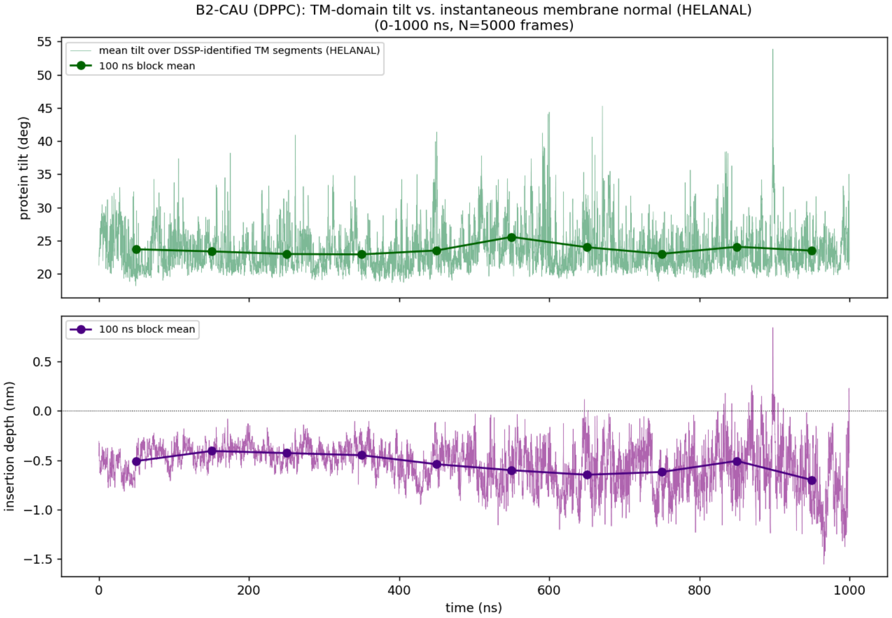

**Figure S3.2.** *Embedding geometry. Transmembrane-domain tilt relative to the instantaneous membrane normal (DSSP-identified TM segments; HELANAL axis for helices, Bansal, Kumar & Velavan 2000, or direct end-to-end C-alpha vector for OmpG's beta strands), and insertion depth of the TM-domain centroid relative to the bilayer midplane, both vs. time with 100 ns block means.*

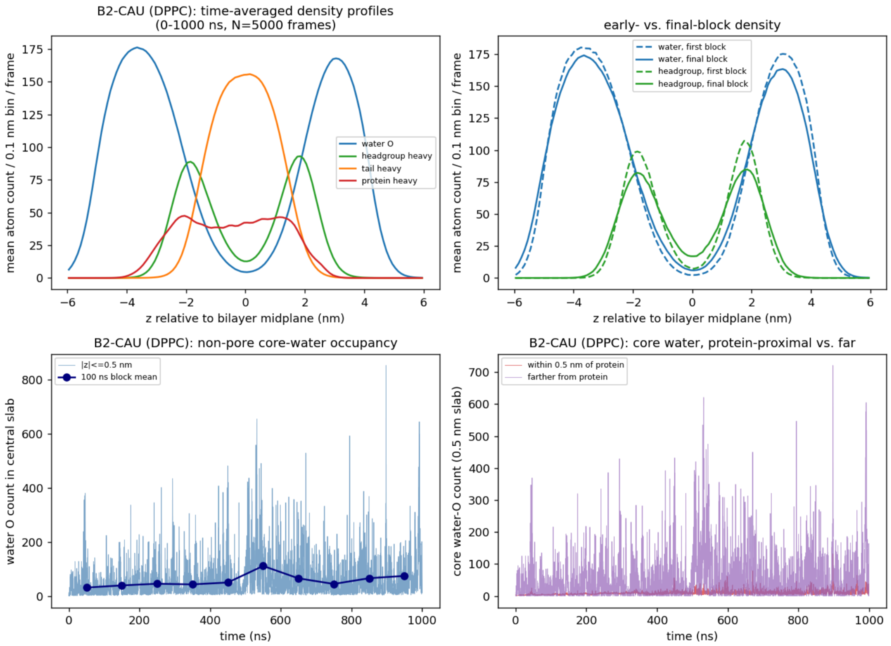


**Figure S3.3.** *Density and water defects. Time-averaged 1D number-density profiles (water, lipid headgroup, lipid tail, protein) along the membrane normal; early- vs. final-block density comparison; central-slab water occupancy vs. time (pore/lumen vs. lipid-matrix split using a HOLE-derived pore radius, Smart et al. 1996, for KcsA and OmpG; non-pore occupancy for the other four systems); core water decomposed by proximity to protein.*

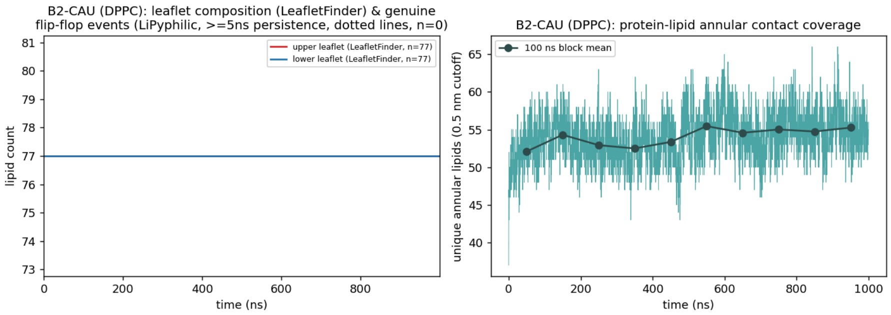


**Figure S3.4.** *Supplementary: leaflet composition (MDAnalysis LeafletFinder counts) with genuine flip-flop events (LiPyphilic, >=5 ns persistence) marked as vertical lines, and unique protein-lipid annular contacts (0.5 nm heavy-atom cutoff) vs. time with 100 ns block means.*

#### D2-GMJ-membrane System

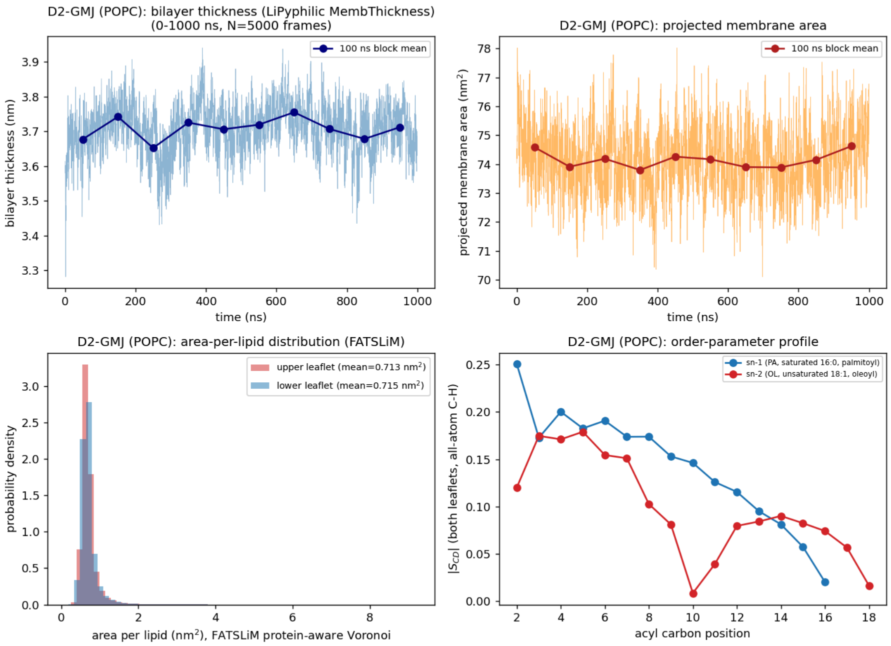


**Figure S4.1.** *Membrane structure. Bilayer thickness and projected membrane area vs. time (raw trace with 100 ns block means); area-per-lipid distribution by leaflet (FATSLiM, protein-aware Voronoi tessellation, Buchoux 2017); mean |S_CD| acyl-chain order-parameter profile for both lipid tails, from explicit C-H bond vectors.*

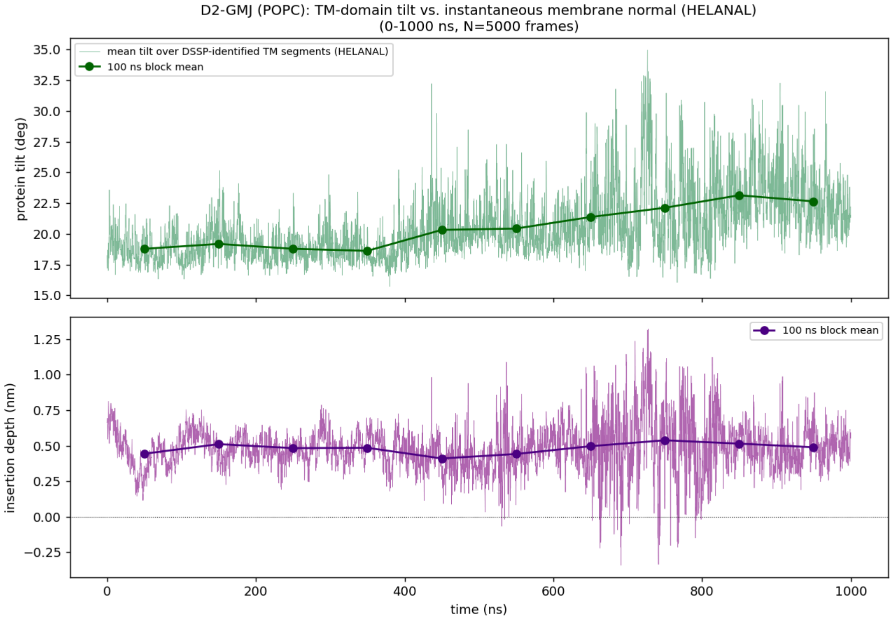

**Figure S4.2.** *Embedding geometry. Transmembrane-domain tilt relative to the instantaneous membrane normal (DSSP-identified TM segments; HELANAL axis for helices, Bansal, Kumar & Velavan 2000, or direct end-to-end C-alpha vector for OmpG's beta strands), and insertion depth of the TM-domain centroid relative to the bilayer midplane, both vs. time with 100 ns block means.*

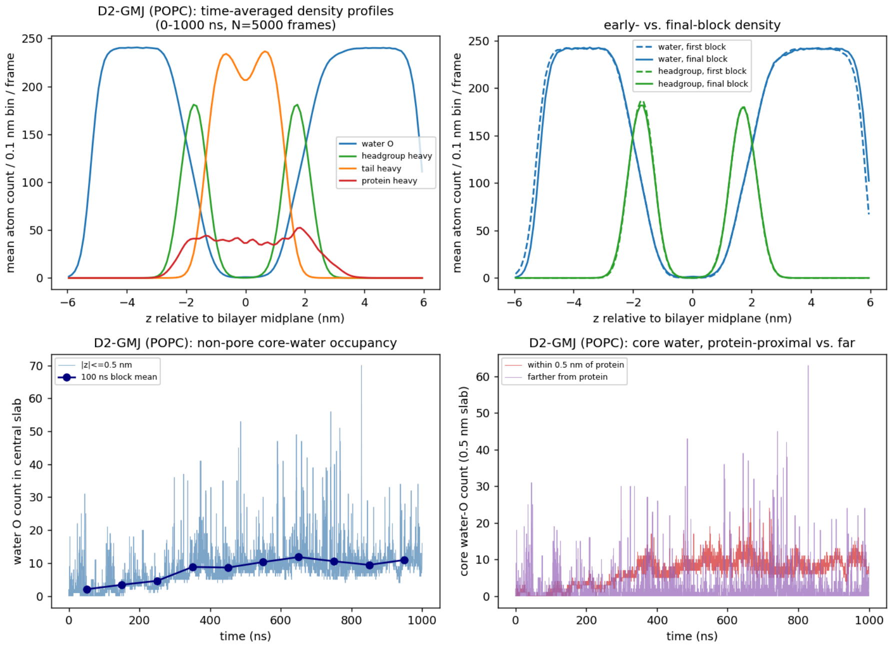


**Figure S4.3.** *Density and water defects. Time-averaged 1D number-density profiles (water, lipid headgroup, lipid tail, protein) along the membrane normal; early- vs. final-block density comparison; central-slab water occupancy vs. time (pore/lumen vs. lipid-matrix split using a HOLE-derived pore radius, Smart et al. 1996, for KcsA and OmpG; non-pore occupancy for the other four systems); core water decomposed by proximity to protein.*

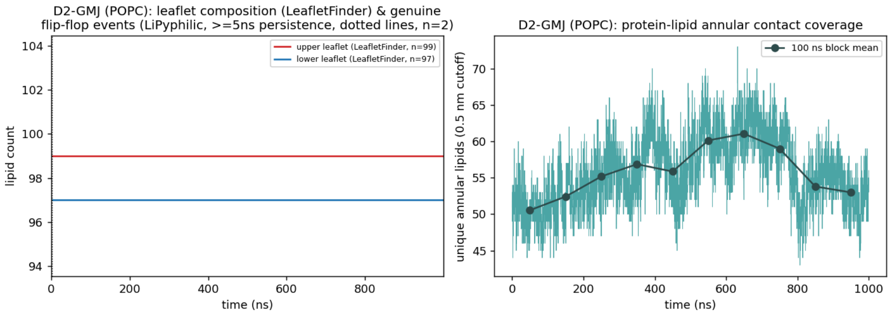


**Figure S4.4.** *Supplementary: leaflet composition (MDAnalysis LeafletFinder counts) with genuine flip-flop events (LiPyphilic, >=5 ns persistence) marked as vertical lines, and unique protein-lipid annular contacts (0.5 nm heavy-atom cutoff) vs. time with 100 ns block means.*

#### KcsA-membrane System

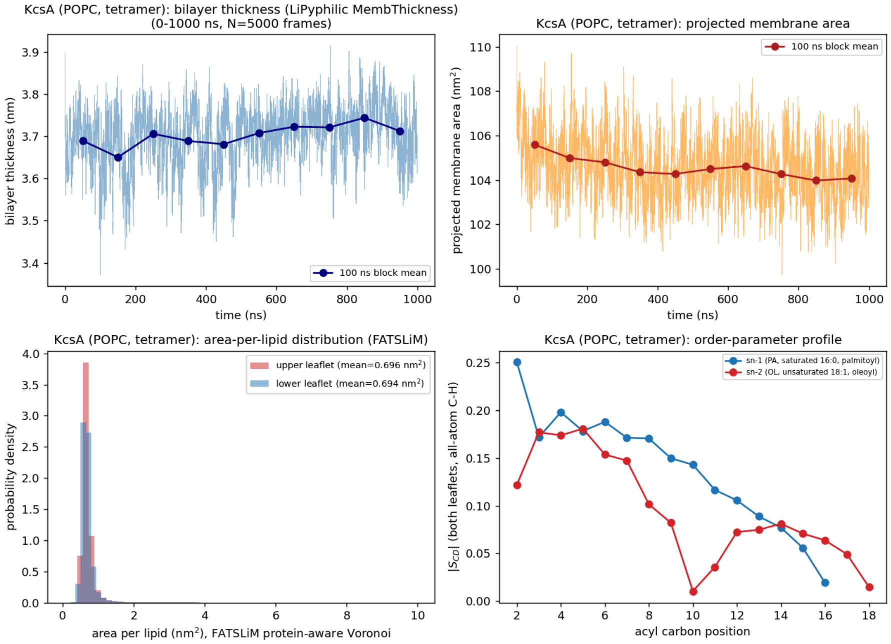


**Figure S5.1.** *Membrane structure. Bilayer thickness and projected membrane area vs. time (raw trace with 100 ns block means); area-per-lipid distribution by leaflet (FATSLiM, protein-aware Voronoi tessellation, Buchoux 2017); mean |S_CD| acyl-chain order-parameter profile for both lipid tails, from explicit C-H bond vectors.*

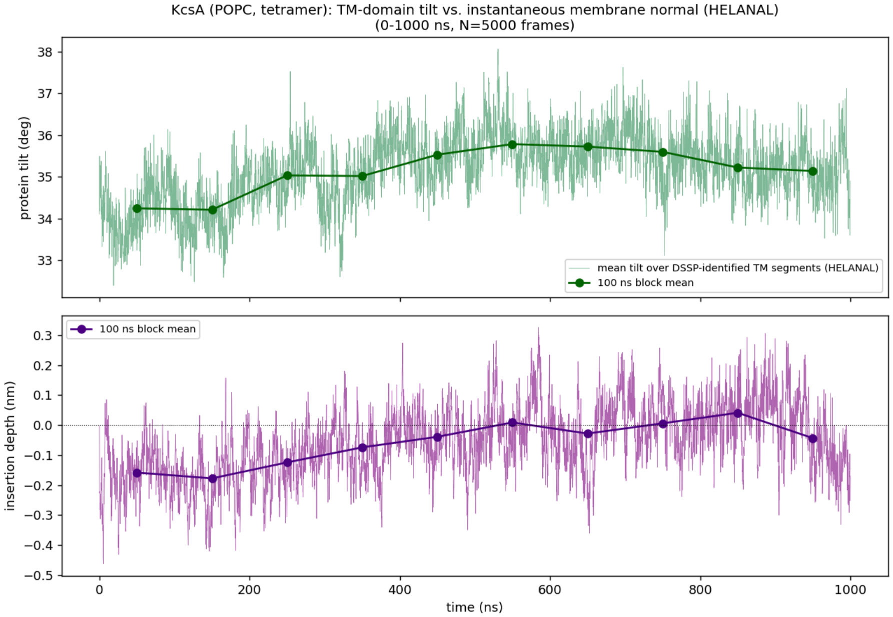

**Figure S5.2.** *Embedding geometry. Transmembrane-domain tilt relative to the instantaneous membrane normal (DSSP-identified TM segments; HELANAL axis for helices, Bansal, Kumar & Velavan 2000, or direct end-to-end C-alpha vector for OmpG's beta strands), and insertion depth of the TM-domain centroid relative to the bilayer midplane, both vs. time with 100 ns block means.*

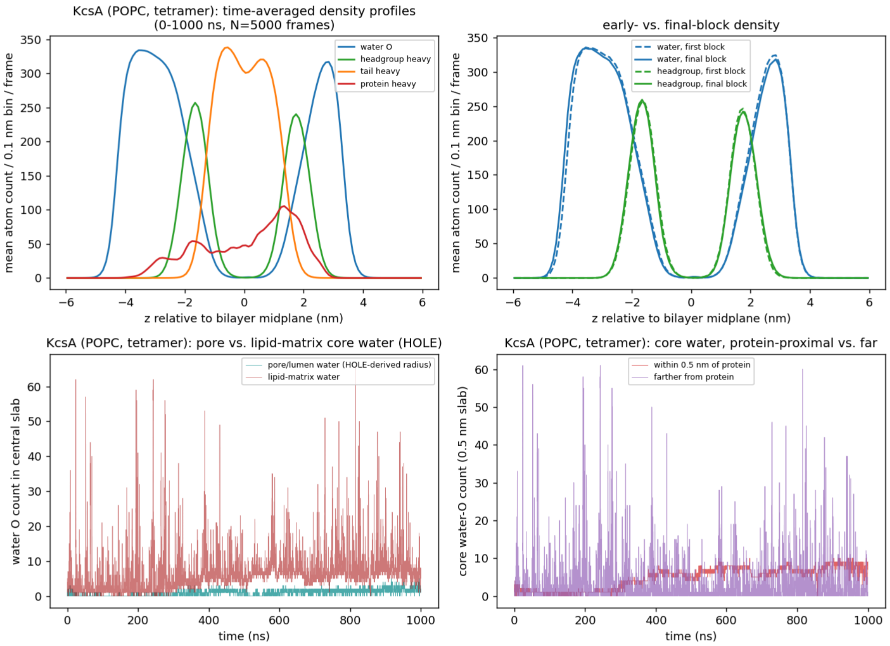


**Figure S5.3.** *Density and water defects. Time-averaged 1D number-density profiles (water, lipid headgroup, lipid tail, protein) along the membrane normal; early- vs. final-block density comparison; central-slab water occupancy vs. time (pore/lumen vs. lipid-matrix split using a HOLE-derived pore radius, Smart et al. 1996, for KcsA and OmpG; non-pore occupancy for the other four systems); core water decomposed by proximity to protein.*

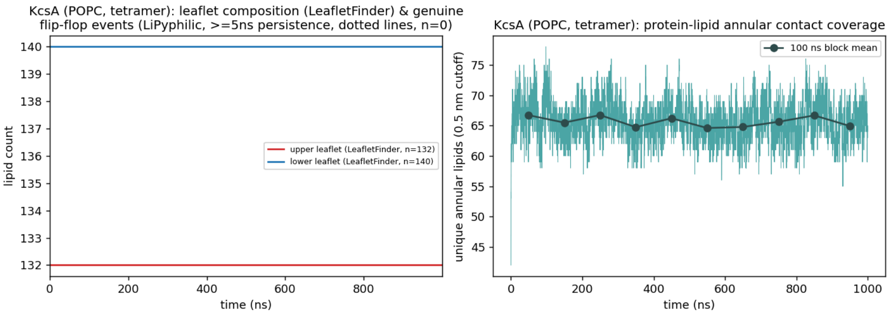


**Figure S5.4.** *Supplementary: leaflet composition (MDAnalysis LeafletFinder counts) with genuine flip-flop events (LiPyphilic, >=5 ns persistence) marked as vertical lines, and unique protein-lipid annular contacts (0.5 nm heavy-atom cutoff) vs. time with 100 ns block means.*

#### M1-CLE-membrane System

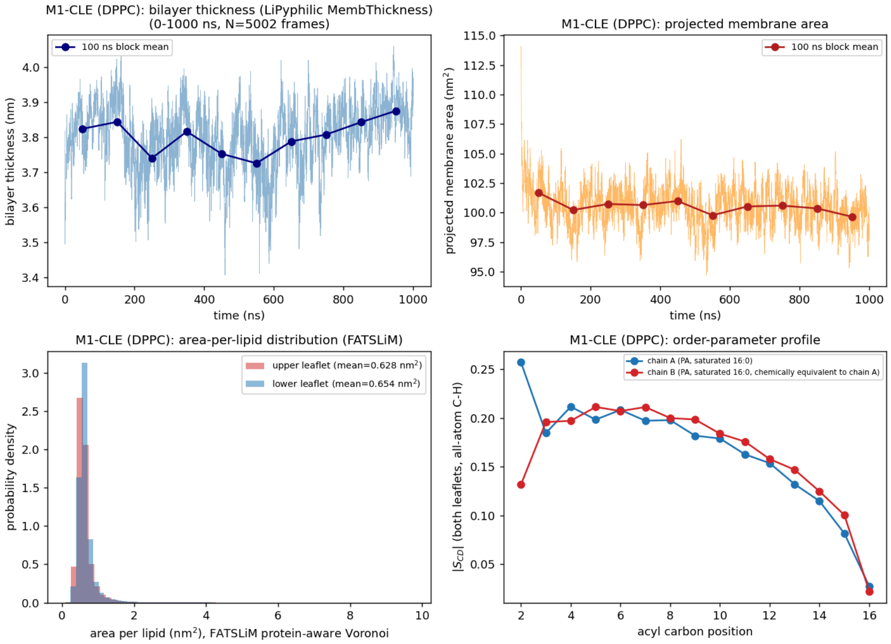


**Figure S6.1.** *Membrane structure. Bilayer thickness and projected membrane area vs. time (raw trace with 100 ns block means); area-per-lipid distribution by leaflet (FATSLiM, protein-aware Voronoi tessellation, Buchoux 2017); mean |S_CD| acyl-chain order-parameter profile for both lipid tails, from explicit C-H bond vectors.*

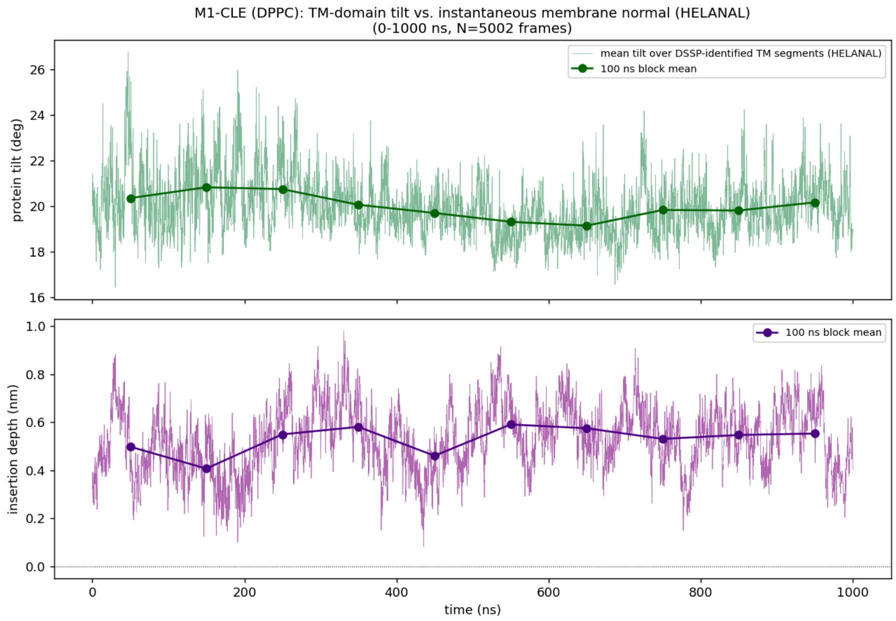


**Figure S6.2.** *Embedding geometry. Transmembrane-domain tilt relative to the instantaneous membrane normal (DSSP-identified TM segments; HELANAL axis for helices, Bansal, Kumar & Velavan 2000, or direct end-to-end C-alpha vector for OmpG's beta strands), and insertion depth of the TM-domain centroid relative to the bilayer midplane, both vs. time with 100 ns block means.*

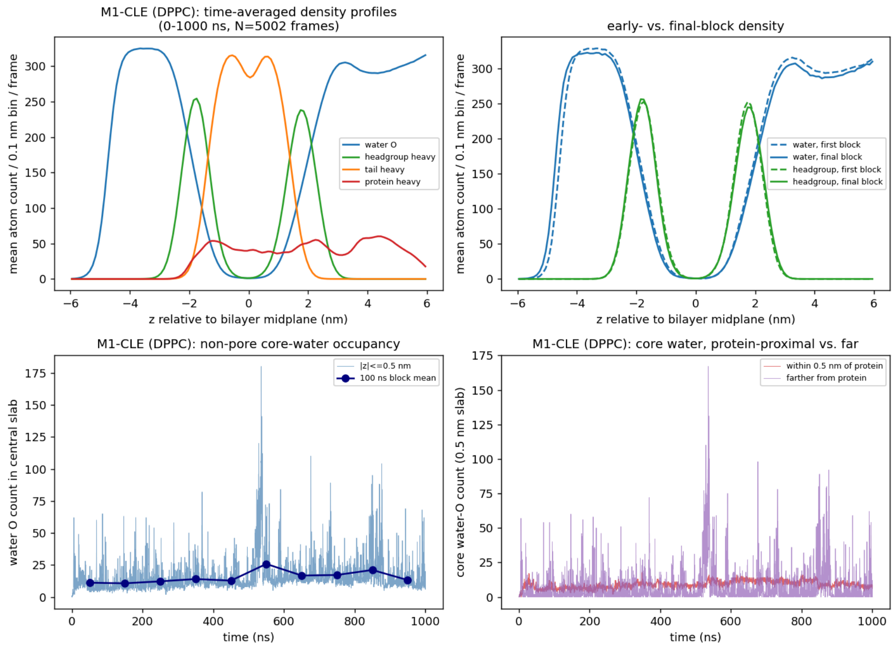


**Figure S6.3.** *Density and water defects. Time-averaged 1D number-density profiles (water, lipid headgroup, lipid tail, protein) along the membrane normal; early- vs. final-block density*

*comparison; central-slab water occupancy vs. time (pore/lumen vs. lipid-matrix split using a HOLE-derived pore radius, Smart et al. 1996, for KcsA and OmpG; non-pore occupancy for the other four systems); core water decomposed by proximity to protein.*

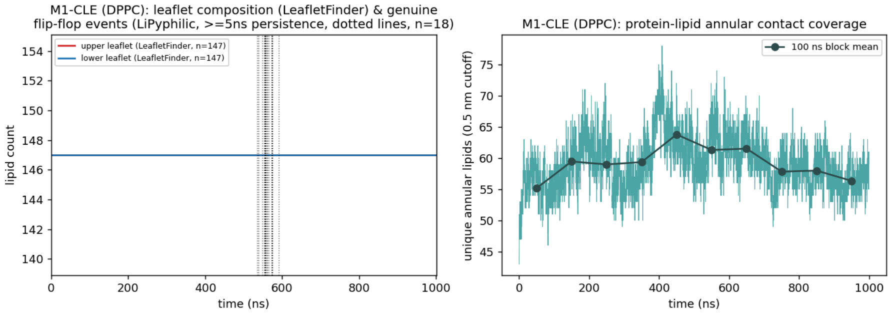


**Figure S6.4.** *Supplementary: leaflet composition (MDAnalysis LeafletFinder counts) with genuine flip-flop events (LiPyphilic, >=5 ns persistence) marked as vertical lines, and unique protein-lipid annular contacts (0.5 nm heavy-atom cutoff) vs. time with 100 ns block means.*

#### M2-IXO-membrane System

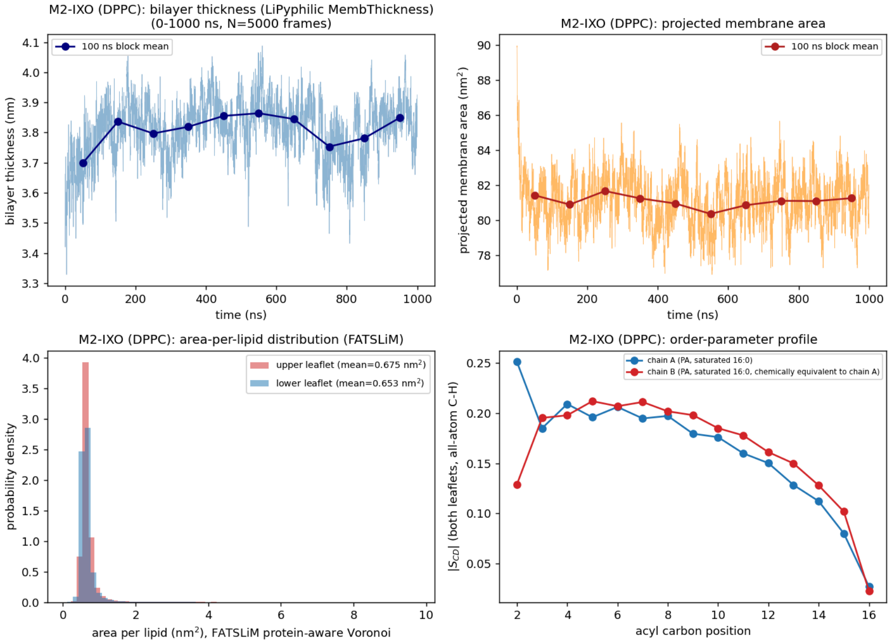


**Figure S7.1.** *Membrane structure. Bilayer thickness and projected membrane area vs. time (raw trace with 100 ns block means); area-per-lipid distribution by leaflet (FATSLiM, protein-aware Voronoi tessellation, Buchoux 2017); mean |S_CD| acyl-chain order-parameter profile for both lipid tails, from explicit C-H bond vectors.*

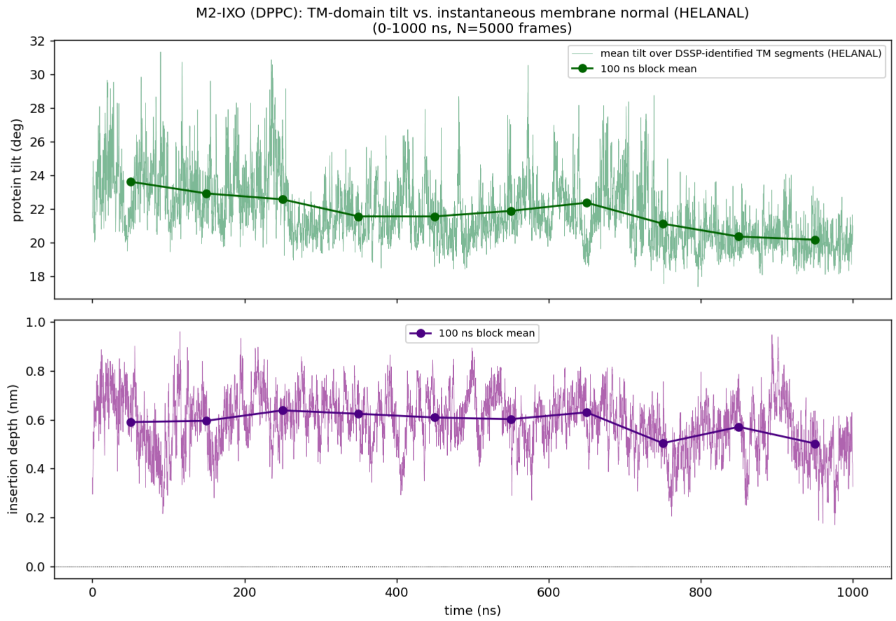


**Figure S7.2.** *Embedding geometry. Transmembrane-domain tilt relative to the instantaneous membrane normal (DSSP-identified TM segments; HELANAL axis for helices, Bansal, Kumar & Velavan 2000, or direct end-to-end C-alpha vector for OmpG's beta strands), and insertion depth of the TM-domain centroid relative to the bilayer midplane, both vs. time with 100 ns block means.*

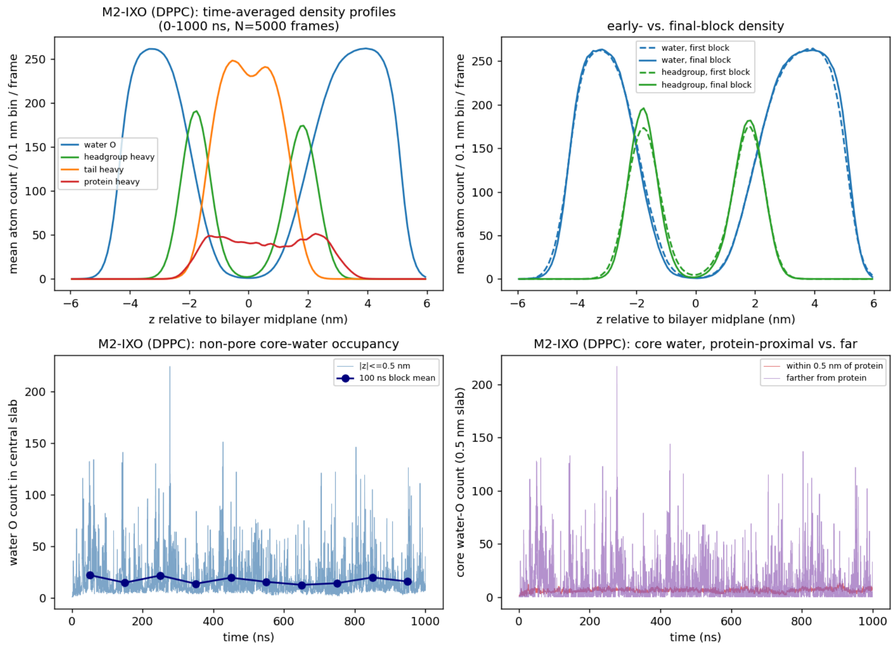


**Figure S7.3.** *Density and water defects. Time-averaged 1D number-density profiles (water, lipid headgroup, lipid tail, protein) along the membrane normal; early- vs. final-block density*

*comparison; central-slab water occupancy vs. time (pore/lumen vs. lipid-matrix split using a HOLE-derived pore radius, Smart et al. 1996, for KcsA and OmpG; non-pore occupancy for the other four systems); core water decomposed by proximity to protein.*

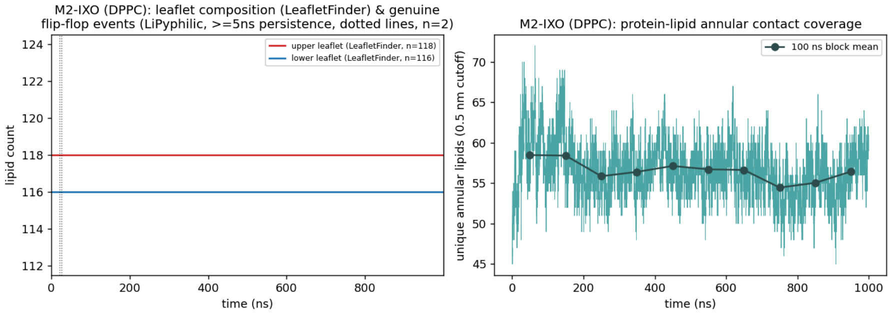


**Figure S7.4.** *Supplementary: leaflet composition (MDAnalysis LeafletFinder counts) with genuine flip-flop events (LiPyphilic, >=5 ns persistence) marked as vertical lines, and unique protein-lipid annular contacts (0.5 nm heavy-atom cutoff) vs. time with 100 ns block means.*

#### OmpG-membrane System

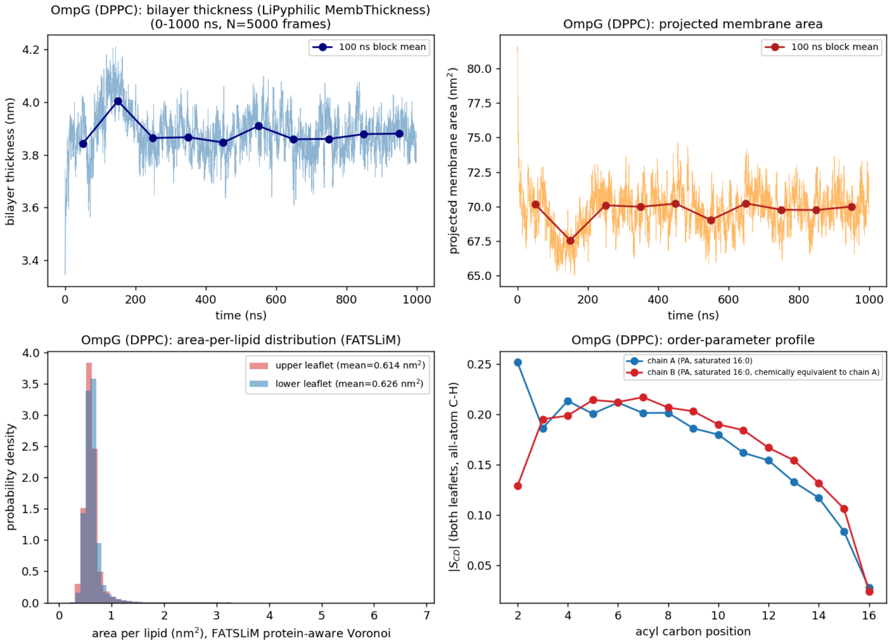


**Figure S8.1.** *Membrane structure. Bilayer thickness and projected membrane area vs. time (raw trace with 100 ns block means); area-per-lipid distribution by leaflet (FATSLiM, protein-aware Voronoi tessellation, Buchoux 2017); mean |S_CD| acyl-chain order-parameter profile for both lipid tails, from explicit C-H bond vectors.*

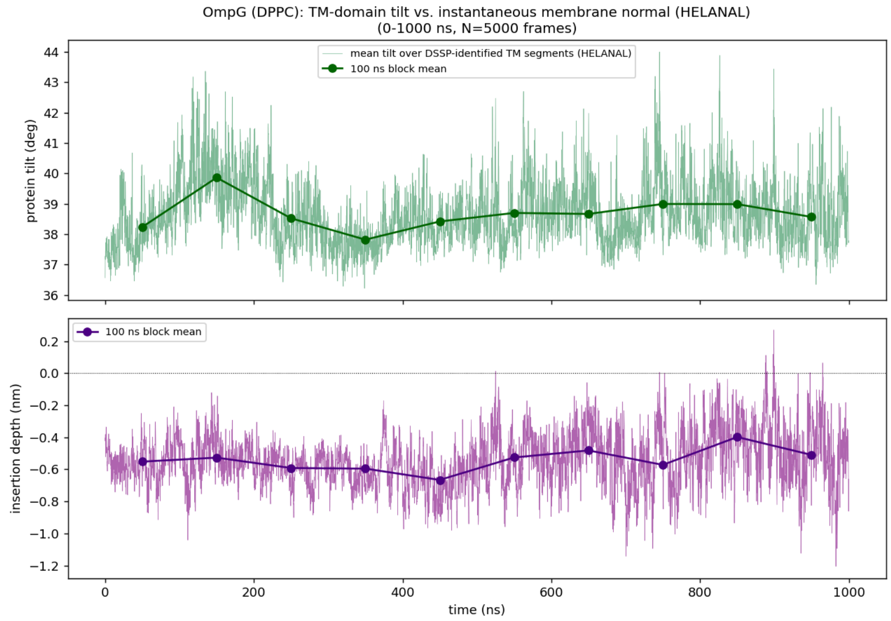


**Figure S8.2.** *Embedding geometry. Transmembrane-domain tilt relative to the instantaneous membrane normal (DSSP-identified TM segments; HELANAL axis for helices, Bansal, Kumar & Velavan 2000, or direct end-to-end C-alpha vector for OmpG's beta strands), and insertion depth of the TM-domain centroid relative to the bilayer midplane, both vs. time with 100 ns block means.*

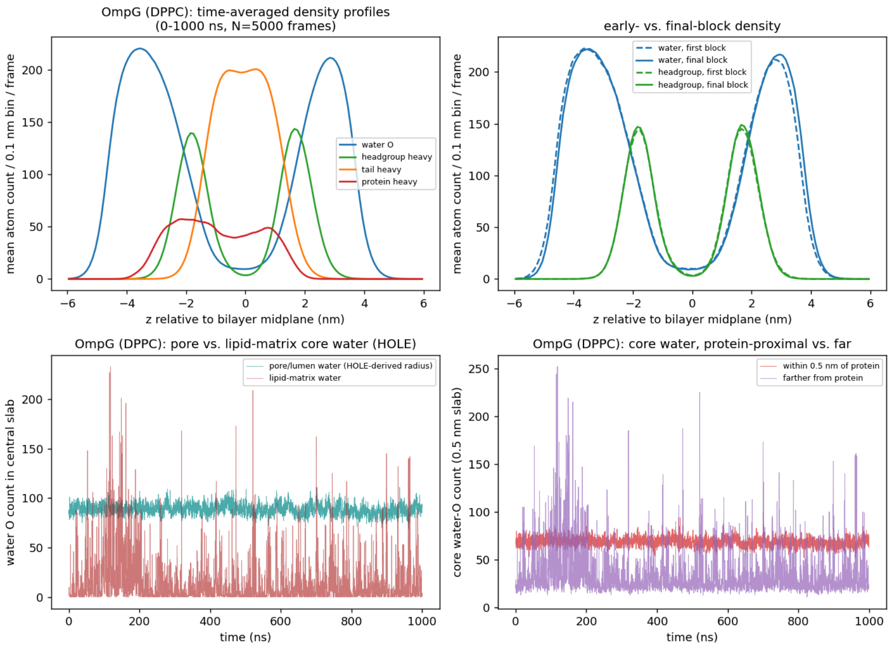


**Figure S8.3.** *Density and water defects. Time-averaged 1D number-density profiles (water, lipid headgroup, lipid tail, protein) along the membrane normal; early- vs. final-block density*

*comparison; central-slab water occupancy vs. time (pore/lumen vs. lipid-matrix split using a HOLE-derived pore radius, Smart et al. 1996, for KcsA and OmpG; non-pore occupancy for the other four systems); core water decomposed by proximity to protein.*

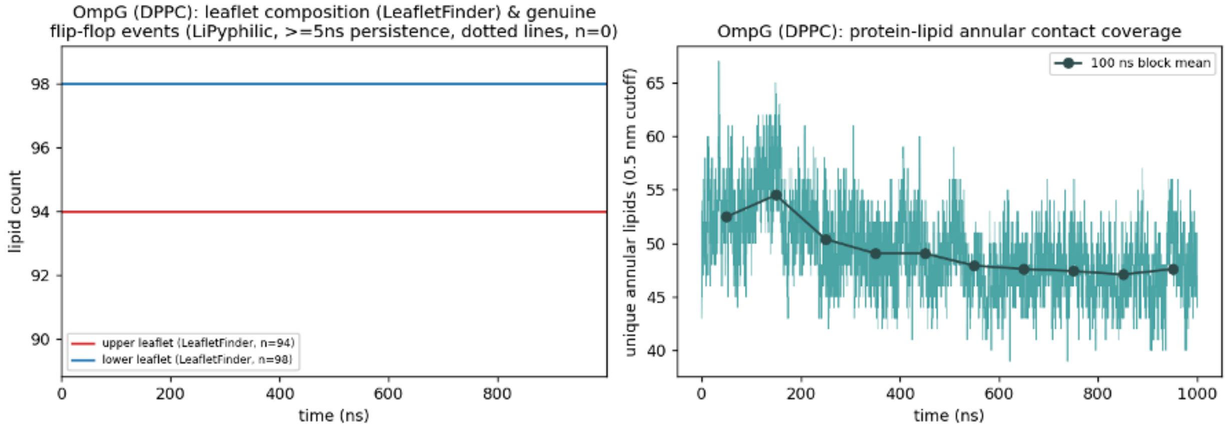


**Figure S8.4.** *Supplementary: leaflet composition (MDAnalysis LeafletFinder counts) with genuine flip-flop events (LiPyphilic, >=5 ns persistence) marked as vertical lines, and unique protein-lipid annular contacts (0.5 nm heavy-atom cutoff) vs. time with 100 ns block means.*
